# A role for selective autophagy of the ER in gametogenic rejuvenation revealed by microfluidics-based lifespan profiling

**DOI:** 10.1101/2025.10.07.681065

**Authors:** Silvan Spiri, Tina Lynn Sing, Nhi Phung, Jay S Goodman, Elçin Ünal, Gloria Ann Brar

**Affiliations:** Department of Molecular and Cell Biology, University of California, Berkeley, Berkeley, CA, 94720, USA; California Institute for Quantitative Biosciences (QB3), University of California, Berkeley, CA, 94720, USA; Center for Computational Biology, University of California, Berkeley, Berkeley, CA 94720, USA

**Keywords:** gametogenesis, rejuvenation, lifespan, microfluidics, autophagy, *Saccharomyces cerevisiae*, meiosis, yeast, ER-phagy

## Abstract

During mitotic growth, *Saccharomyces cerevisiae* cells age by dividing asymmetrically producing young daughter cells while retaining age-associated damage in the mother cell, which will eventually become senescent. Gametogenesis naturally and fully resets precursor cell lifespan, even for replicatively aged cells. However, the mechanisms responsible for gametogenic rejuvenation remain elusive. This is, in part, due to the existing methods to quantify replicative lifespan resetting in this context, which are limited to low-throughput and labor-intensive approaches. Here, we introduce a high-throughput microfluidic-based assay that allows systematic characterization of factors required for gametogenic rejuvenation in *S. cerevisiae.* With this technique, we show that we can sensitively measure a wide range of gamete replicative lifespans that are consistent with known short and long-lived mutants. Excitingly, using this technique, we report Atg39 and Atg40, receptors involved in selective autophagy of the ER, as the first identified molecular determinants of gametogenic rejuvenation. We anticipate that this novel technique will enable systematic identification of additional molecular factors that drive gametogenic rejuvenation.

## Introduction

Life is continuously maintained by organisms through the genesis of offspring. In sexually reproducing eukaryotes, gametogenesis is a conserved developmental program that forms haploid gametes from diploid precursor cells. Although somatic cells deteriorate over time, robust mechanisms eliminate age-associated damage in the germline and during gametogenesis to secure the fitness and survival of the next generation (Gladyshev, 2021; Jones, 2007; Maklakov and Immler, 2016; Sing et al., 2022; Smelick and Ahmed, 2005; Xiao and Ünal, 2025; Yamashita, 2023). Despite the importance of these pathways, the specific molecular mechanisms required for mediating this programmed developmental rejuvenation remain elusive.

During mitotic growth, *Saccharomyces cerevisiae* divides asymmetrically, producing young daughter cells and retaining age-associated damage in the aging mother cell (Denoth Lippuner et al., 2014; Higuchi-Sanabria et al., 2014; Hughes and Gottschling, 2012; Nyström and Liu, 2014). This limits the number of daughter cells that a mother cell can produce (termed “replicative lifespan”) before reaching senescence. Excitingly, upon completion of gametogenesis, replicative lifespan of gametes is completely reset in *S. cerevisiae*, relative to that of the precursor cell (Sing et al., 2022; Ünal et al., 2011; Ünal and Amon, 2011) (**Fig. 1**). The portability of this rejuvenation program was demonstrated by the observation that ectopic expression of Ndt80, a transcription factor necessary for completion of gametogenesis, extends replicative lifespan in mitotic cells (Ünal et al., 2011; Xu et al., 1995). Since this initial discovery, there has been interest in determining the gametogenic factors that drive lifespan resetting in gametes (Sing et al., 2022). Drug-mediated inhibition of bulk autophagy prevents clearance of age-associated Hsp104 foci during gametogenesis as well as the completion of gametogenesis. This suggests that autophagy-mediated turnover of aged proteins and cellular components may play an important role in orchestrating gametogenic rejuvenation (Tyler and Johnson, 2018; Ünal et al., 2011).

**Figure 1.**
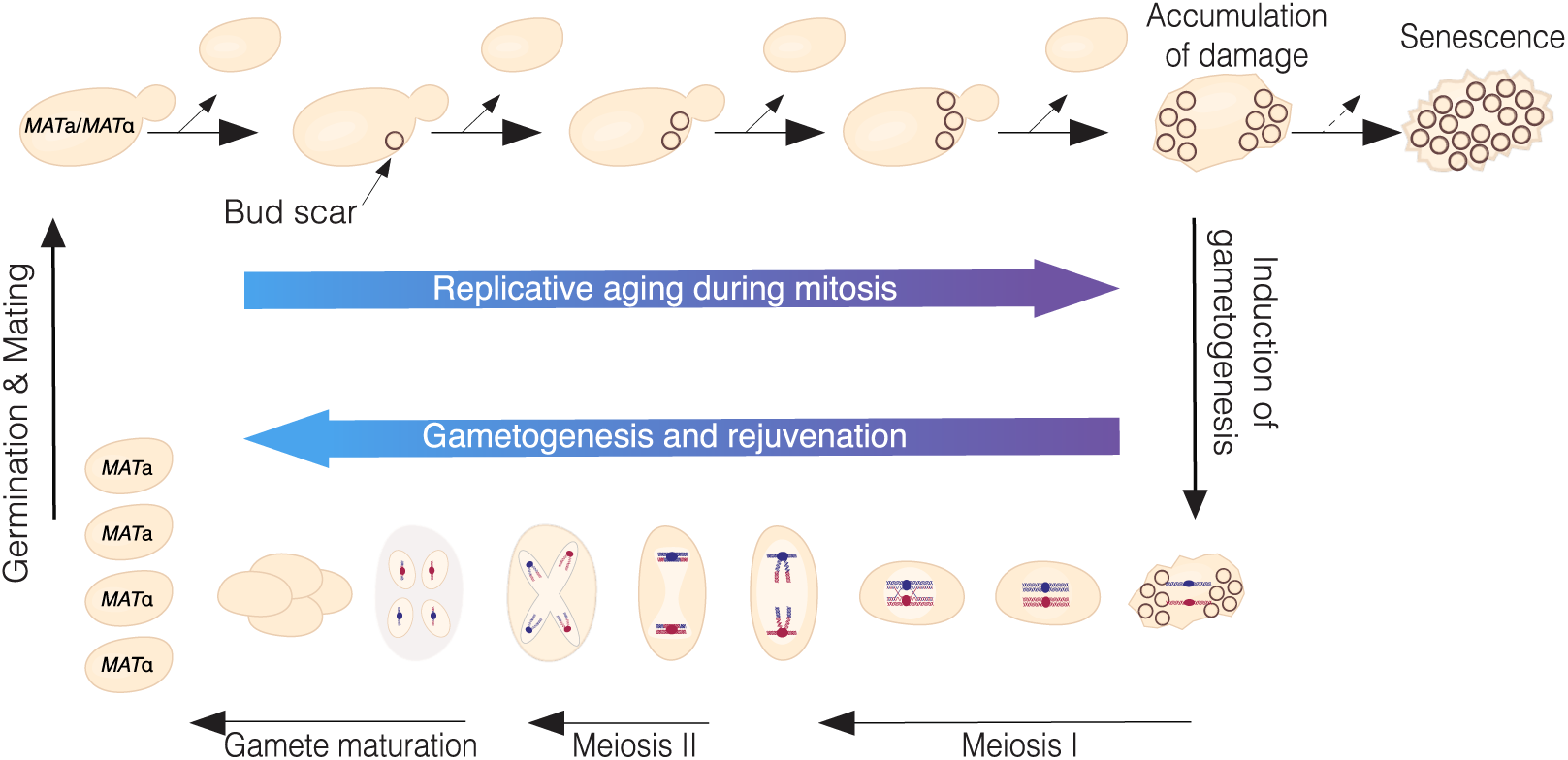
Replicative lifecycle and gametogenic rejuvenation in *Saccharomyces cerevisiae*. In vegetative growth, yeast cells divide asymmetrically, with the daughter cell born young while the mother cell accumulates aged and damaged components until reaching senescence. Replicative age of a single yeast cell is determined by the number of bud scars, which form at each division and are retained on the cell wall (top panel). Gametogenesis can be induced by nutrient depletion (downwards arrow, right). During gametogenesis, diploid yeast divides symmetrically, fully resetting their lifespan and leading to the formation of four equally young haploid gametes with two different mating types, even from aged precursor cells (bottom panel). Addition of nutrients induces germination, return to mitotic growth and formation of diploid cells upon mating (upward arrow, left).

Yeast gametogenesis, a highly experimentally tractable program that can be induced efficiently at will, presents the unique opportunity to gain mechanistic insights into the process of gametogenic rejuvenation. However, current techniques to quantify replicative lifespan of gametes significantly limit progress in the field. Quantification of gamete replicative lifespan has relied on scoring the replicative lifespan of a gamete over several days by continuous counting and manual removal of daughter cells via micromanipulation (Ünal et al., 2011). This technique is tedious, low throughput and labor-intensive. In fact, since gametogenesis-specific rejuvenation was initially observed (Ünal et al., 2011), only two other studies have used this technique to measure replicative lifespan of gametes (Koch et al., 2020; Lee et al., 2019) and the specific pathways required for directly mediating gametogenic rejuvenation remain unclear. Although micromanipulation has historically been used to quantify replicative lifespan in vegetatively growing haploid yeast cells (Mortimer and Johnston, 1959), new microfluidics-based approaches that increase throughput and lower manual labor have successfully been developed and utilized for yeast lifespan measurement in recent years. Multiple different trap designs have been used to capture and retain mother cells in imaging chambers with a constant media flow over prolonged periods of time. Parallelization of multiple imaging chambers allows acquisition of thousands of cells from multiple different genetic backgrounds over a few days in one single experiment (Chen et al., 2017; Jin et al., 2019; Jo et al., 2015; Li et al., 2020; Liu et al., 2015; Thayer et al., 2022). While this method is rapidly replacing micromanipulation-based readouts for vegetatively growing haploid yeast cells, a microfluidics-based platform to investigate gamete replicative lifespan does not currently exist. The challenges associated with capturing and measuring gamete lifespan include: (i) purifying gametes from precursor cells and from the ascus that encloses four gametes together, (ii) preventing mating of neighboring gametes, and (iii) the small size of gametes relative to precursor cells.

To overcome these limitations, we developed a microfluidic-based microscopy assay that allows high-throughput characterization of factors required for gametogenic rejuvenation in *S. cerevisiae*. We engineered inert genetic modifications and mitigated the aforementioned challenges to enable microfluidic-based assessment of replicative lifespan in *S. cerevisiae* gametes derived from the SK1 background, which undergoes gametogenesis with high efficiency (Kane and Roth, 1974; Padmore et al., 1991). With this approach, we were able to isolate and purify single gametes from both young and aged precursor cells, which were then imaged over 72 hours using a custom microfluidic device, enabling quantitative analysis of gamete replicative lifespan.

Comparing the replicative lifespan of gametes derived from young versus aged precursor cells captured a broad spectrum of lifespan outcomes and confirmed the effects of known short- and long-lived mutants. Most excitingly, using this system, we discovered that turnover of the endoplasmic reticulum (ER) mediated by selective autophagy (ER-phagy) but not mitophagy is required during gametogenesis for complete lifespan resetting in gametes derived from aged precursor cells. ER-phagy receptors Atg39 and Atg40 represent the first proteins known to drive full lifespan resetting through gametogenesis, providing a molecular foothold to dissect the impact of ER quality and inheritance in cellular deterioration. Moreover, this discovery exemplifies the potential of our microfluidic approach to uncover the mechanisms by which eukaryotic cells counteract age-associated damage.

## Results

### An approach for long-term marking of gametes to enable gamete lifespan analysis

To microscopically distinguish gametes from precursor cells, we looked for markers exclusively expressed in gametes. One such gene, *PMA2,* encodes a plasma membrane proton pump (Carmelo et al., 1997; Saliba et al., 2018; Serrano et al., 1986). Its paralogous protein, Pma1, is highly expressed during vegetative growth, with an exceptionally long half-life and asymmetric retention in mother cells (Henderson et al., 2014). *PMA1* is essential for viability but *PMA2* is not, and is barely detectable during mitotic growth (Fernandes and Sá-Correia, 2003; Schlesser et al., 1988; Serrano et al., 1986). *PMA2* translation and protein levels are upregulated late in gametogenesis (**Fig. S1A-C**) (Cheng et al., 2019). To further investigate the expression pattern of *PMA2*, we created a transgenic reporter strain carrying a Pma2-GFP fusion protein under control of its endogenous promoter and tracked its expression timing relative to meiotic divisions by using a fluorescently labeled chromatin marker (Htb1-mCherry). Pma2-GFP expression was upregulated 1-2 hours following anaphase II and predominantly localized to the gamete plasma membranes (**Fig. 2A**). Importantly, Pma2-GFP was not detected in cells that fail to undergo gametogenesis (arrows, **Fig. 2A**). Next, we analyzed Pma2-GFP expression in gametes undergoing germination, their first mitotic division. We found that gamete-derived daughter cells lacked Pma2-GFP signal, which was fully retained on the plasma membrane of gametes for several rounds of cell division (**Fig. 2B**). This expression pattern analysis revealed that Pma2-GFP is a reliable, specific, and retained marker of gametes exhibiting gamete-specific retention beyond germination, making it an optimal reporter to distinguish gametes cells that fail to complete gametogenesis and vegetative daughter cells.

**Figure 2.**
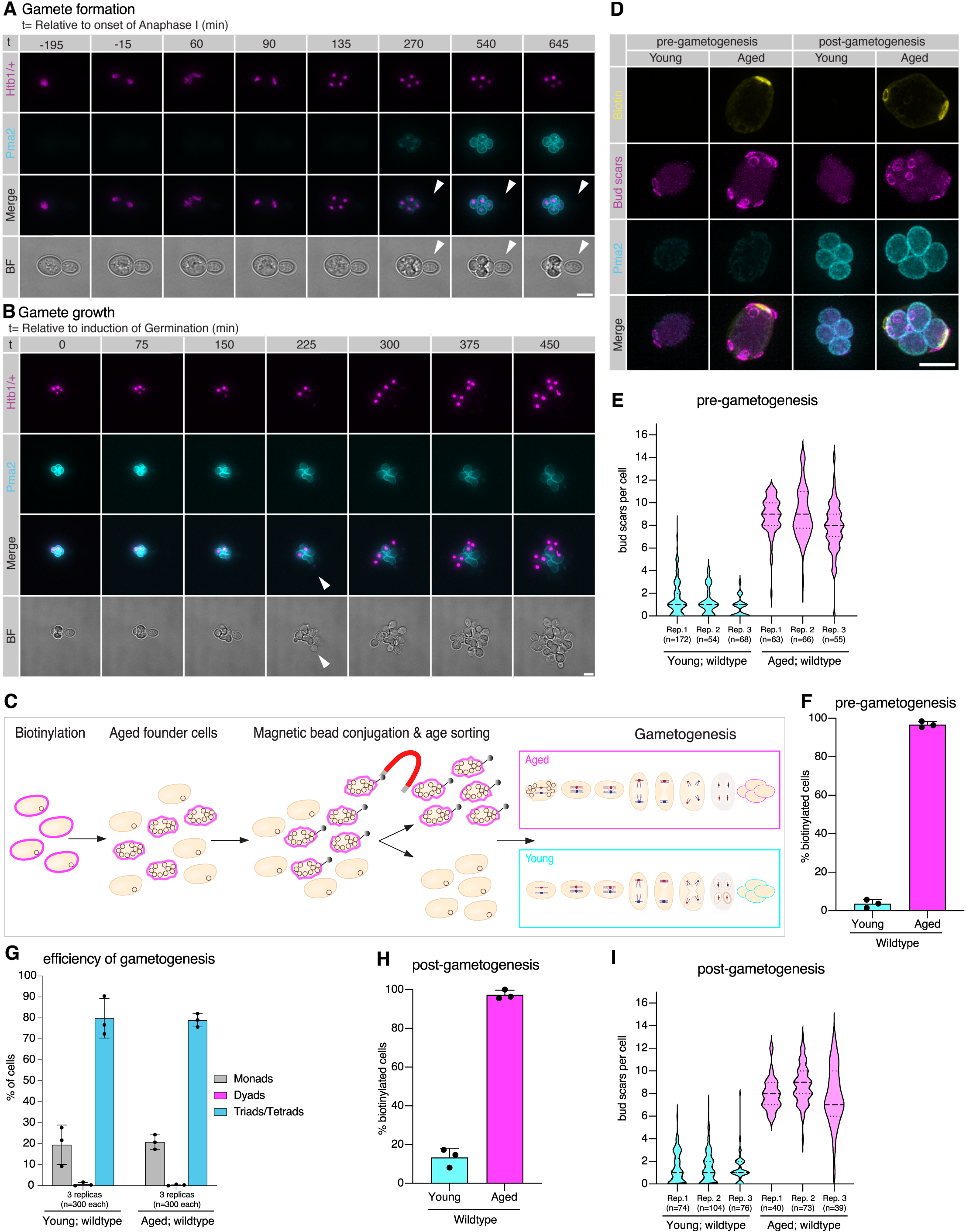
Gametogenesis-specific expression of Pma2 remains detectable post-germination in age-sorted gametes. (**A**) Montage of max-intensity projected diploid cell progressing through gametogenesis visualizing Pma2-GFP (cyan) and a Htb1-mCherry (magenta) marking histones, arrows indicate a cell that failed to go through gametogenesis. (**B**) Montage of a germinating max-intensity projected tetrad visualizing Pma2-GFP (cyan) and a Htb1-mCherry (magenta) marking histones, arrow indicates Pma2-GFP negative emerging daughter cell. (**C**) Schematic depiction of age sorting by biotin mediated magnetic bead conjugation. (**D**) Representative age sorted pre-gametogenic max-intensity projected cells (left panel) and post-gametogenic tetrads (right panel) visualizing biotin label (yellow), chitin rich bud scars (magenta) and Pma2-GFP (cyan). (**E**) Quantification of replicative age by assessing number of bud scars in young (1.16 ± 0.41, mean ± SD, n=3 replicas, n=172; 54 and 68 respectively) and aged (8.55 ± 0.66 (mean ± SD, 3 replicas, n=63, 66 and 55 respectively) wildtype cell populations. (**F**) Quantification of fraction of biotinylated cells in young (3.63 ± 2.12 %, mean ± SD, 3 replicas, n=175, 54 and 68 respectively) and aged (96.74 ± 1.58 %, mean ± SD, 3 replicas, n=65, 66 and 55 respectively) wildtype cell populations. (**G**) Quantification of fraction of tetrads in young (79.8 ± 9.43%; mean ± SD, 3 replicas n=300 each) and aged (aged: 78.89 ± 3.17 %, mean ± SD, 3 replicas n=300 each) post-gametogenic wildtype cell populations. (**H**) Quantification of fraction of biotinylated tetrads in young (13.38 ± 4.74%, mean ± SD, n=3 replicas, n=74, 104 and 76 respectively) and aged (97.34 ± 2.33%, mean ± SD, 3 replicas, n=46, 73 and 39 respectively) wildtype populations after completion of gametogenesis. (**I**) Quantification of replicative age by assessing number of bud scars in young (1.31 ± 0.04, mean ± SD, 3 replicas, n=74, 104 and 76 respectively) and aged (8.17 ± 0.54, mean ± SD, 3 replicas, n=40, 73 and 39 respectively) wildtype tetrad populations. Scale bars are 5 µm and confidence intervals were calculated using the Wilson score method.

### Cell populations efficiently sorted by age progress robustly through gametogenesis

To be able to quantify gametogenic rejuvenation, we first separated pre-meiotic diploid cell populations by replicative age using an established protocol (Smeal et al., 1996). Briefly, a founder cell population was biotin-labeled with sulfo-NHS-LC-biotin, which specifically and permanently binds to cell surface proteins that are retained on the founder (mother) cells during subsequent mitotic divisions. These biotin-labeled founder cells were then replicatively aged and incubated with anti-biotin magnetic beads that enabled their isolation from their young daughter cells. The labeled cell suspension was sequentially passed through two magnetic columns to increase purity, followed by elution. Flow-through cells from initial magnetic sorting provides a pure population of young cells. The eluate from the second column represents the aged cell population (**Fig. 2C**, see methods). Sorted cell populations were stained with fluorescent dyes visualizing both biotin and the chitin-rich, ring-like structures called bud scars on the yeast cell wall, each of which represents a budding event and thus reveals the replicative age of a cell (Cabib and Bowers, 1971). We measured the purity and age of the sorted population by fluorescent microscopy of these markers (**Fig. 2D**, left panel). With our consecutive-sort protocol, we consistently obtained near pure young (3.63% biotinylated cells with a mean replicative age of 1.16), and aged (96.74% biotinylated cells with a mean replicative age of 8.55) cell populations (**Fig. 2E&F**). Next, we induced gametogenesis in the separated cell populations and analyzed efficiency of gametogenesis after 24 hours. We found that 79.8% of young cells and 78.89% of aged cells successfully formed visibly distinct tetrads (**Fig. 2G**) and that gametes from young and aged cell population both robustly expressed Pma2-GFP after completion of gametogenesis (**Fig. 2D**, right panel). Furthermore, we re-evaluated the purity as well as the age of the sorted population following gametogenesis and confirmed that the relative distributions of young and aged populations were maintained (**Fig. 2H&I**). In summary, we were able to reproducibly isolate young and aged cell populations with high purity and observed no negative effect of our experimental regime on the efficiency of gametogenesis.

### Isolation of single gametes for microfluidics-based replicative lifespan profiling

To be able to capture replicative lifespan of yeast gametes with microfluidic devices, we had to overcome four major obstacles. First, without inert mutations to disrupt associations between individual cells, they were not suitable for microfluidics-based applications. Second, gametogenesis produces gametes with two mating types, thus we had to prevent initiation of mating in response to mating pheromones, which leads to growth arrest and abnormal cell morphology. Third, gametes are packaged into tetrad structures inside an enclosed ascus and do not separate during germination (**Fig. 2B**), rendering assessment of replicative lifespan impossible. Thus, we had to find a way to separate tetrads into single gametes before loading into the microfluidics imaging chambers. Fourth, separated gametes are significantly smaller than vegetative growing cells, and do not fit into the custom-built U-traps in our microfluidics devices that are built to capture vegetatively growing haploid cells (**Fig S2A-D** adapted from (Jo et al., 2015)).

Cells from the SK1 background are known to form both covalent (mother-daughter cell adherence) and non-covalent (flocculation) interactions, which present an issue when using microfluidics-based application due to elevated clogging. To mitigate this, we introduced an inert point mutation into *AMN1* to promote efficient mother-daughter cell separation (Fang et al., 2018). We also deleted *FLO8*, a transcription factor required for flocculation (Kobayashi et al., 1996) to prevent the cells from clumping together. During gametogenesis, a diploid yeast cell undergoes two meiotic divisions forming a tightly packed tetrad containing four haploid gametes with a 2:2 segregation of the two mating types *MAT***a** and *MAT***α** (**Fig. 1&3A**). To avoid triggering the mating response in gametes, we replaced the endogenous promoter of the mating response signaling scaffold gene *STE5* (Chol et al., 1994), with an anhydrotetracycline-inducible promoter (**Fig. S2E**) (Bellí et al., 1998). We confirmed sterility and inducible fertility by mixing *MAT***a** and *MAT***α** cells on control and anhydrotetracycline-containing plates and scoring formation of mated zygotes (**Fig. S2F**).

To obtain single gametes and to eliminate cells that did not proceed through gametogenesis, we incubated post-gametogenesis cultures with zymolyase. This enzyme hydrolyzes the main structural polysaccharide in the cell wall, thus lysing non-gametes and weakening the ascus protecting the four gametes within a tetrad (**Fig. 3A**). Zymolyase-treated tetrads were then gently sonicated to disassociate them into single gametes, and presence of the Pma2-GFP marker in single gametes was confirmed by fluorescence microscopy (**Fig. 3A&B**).

**Figure 3.**
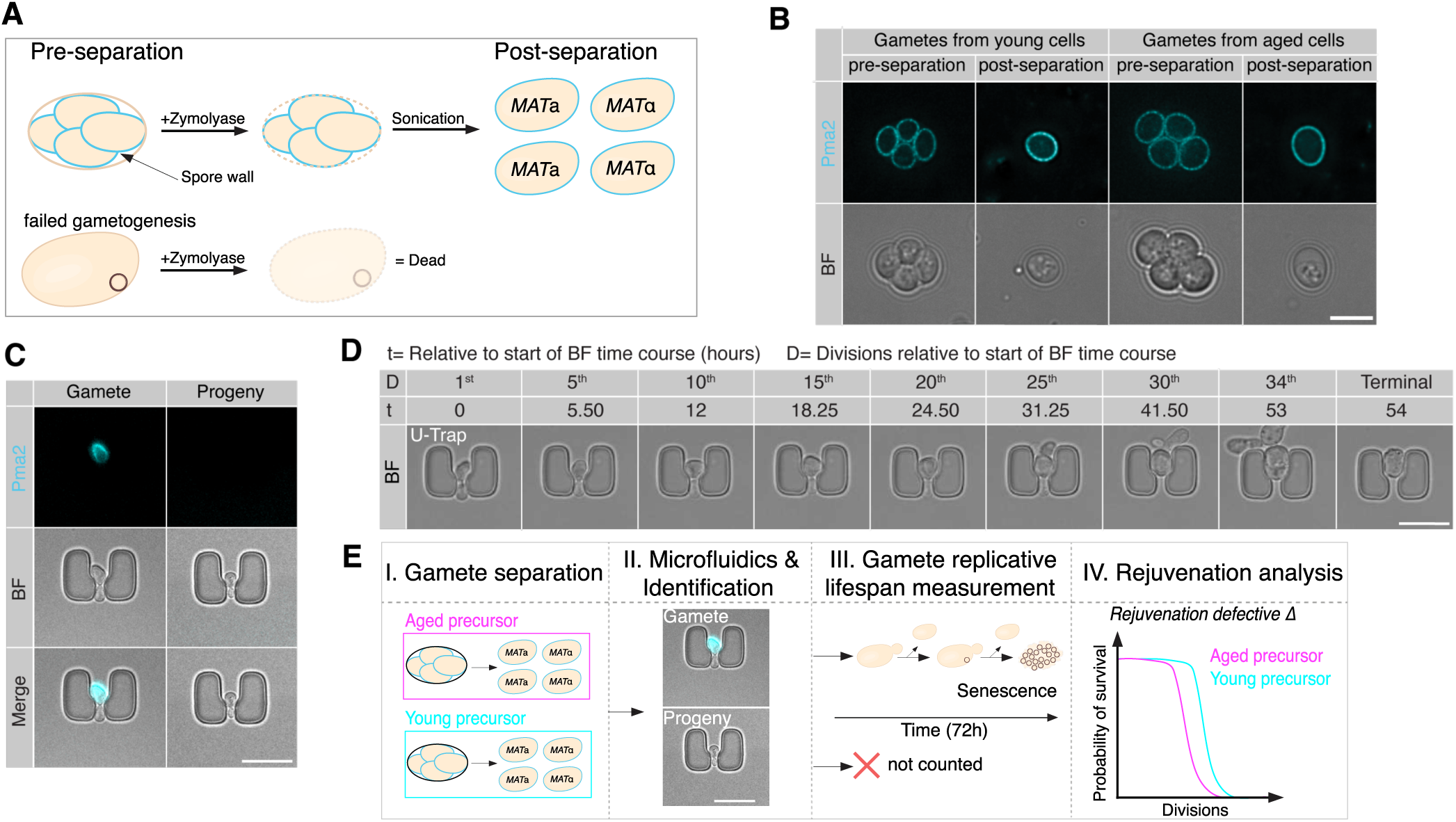
Separation of tetrads into single sterile gametes to assess gamete replicative lifespan with microfluidics. (**A**) Schematic depiction of tetrad separation procedure. (**B**) Representative single z-slices of pre- and post-separated young (left panel) and aged (right panel) tetrads and gametes, respectively, visualizing brightfield and Pma2-GFP (cyan). (**C**) Representative Pma2-GFP positive gamete (left panel) and Pma2-GFP negative gamete progeny (right panel) trapped in microfluidics-based devices visualizing brightfield and Pma2-GFP (cyan). (**D**) Representative gamete trapped in microfluidics imaging unit visualizing every 5^th^ division until the terminal 34^th^ division over the time of 54 hours. (**E**) Schematic depiction of the developed pipeline to assess gametes replicative lifespan. Scale bars are 5 µm.

When loading separated gametes directly into our custom microfluidics devices (**Fig S2A-D** adapted from (Jo et al., 2015)), gametes were not retained in the traps due to their small size. To solve this issue, gametes were first grown in rich media following their separation for 3 hours, which was sufficient to increase their size but only allowed the first cell division to occur (**Fig. S2G**). These pre-grown gametes were then loaded into the microfluidic devices and an image of the whole region of interest was taken in the FITC and brightfield channel. By assessing Pma2-GFP expression, we could reliably determine if a trapped cell was a gamete, or a progeny cell generated during re-growth (**Fig. 3C**). Next to quantify replicative lifespan of gametes, brightfield images were acquired every 15 minutes over 72 hours, capturing every cell division (**Fig. 3D&S2H**). Together, these steps constitute a high-throughput microfluidics-based pipeline that enables for the assessment of gamete replicative lifespan, and identification of mutants that specifically impact the rejuvenation of aged cells (**Fig. 3E**).

### Wild-type gametes from aged precursor cells display complete resetting of replicative lifespan

It has previously been shown that replicative lifespan is completely reset after gametogenesis (Ünal et al., 2011). This study used a micromanipulator to continuously count and remove daughter cells of gametes previously placed onto agar plates and was assessed only in the W303 and A364a strain background. Thus, we wanted to test if gametes also showed a complete reset of their replicative lifespan in our strain background, using the developed microfluidics-based pipeline. Toward this end, we sorted diploid wild-type cells following our newly established protocol (**Fig. 2C**) and analyzed the replicative age of young and aged pre-gametogenic cell populations (**Fig. S3A&B**). Separated cell populations were transferred to gametogenesis-inducing media and after 24 hours both populations successfully completed gametogenesis (**Fig. S3C**). Resulting in pure post-gametogenic tetrad populations from replicatively aged and young precursor cells (**Fig. 4A&S3D**). We acquired and quantified the replicative lifespan of gametes following our pipeline and confirmed that the median replicative lifespan of gametes from both young and aged precursors cells was the same (**Fig. 4B**, p=0.982, Log-rank (Mantel-cox)). An independent biological replicate led to a similar outcome (**Fig. S3E-J**). These results confirm that our newly developed high-throughput microfluidic-based pipeline can capture and quantify gametogenic rejuvenation.

**Figure 4.**
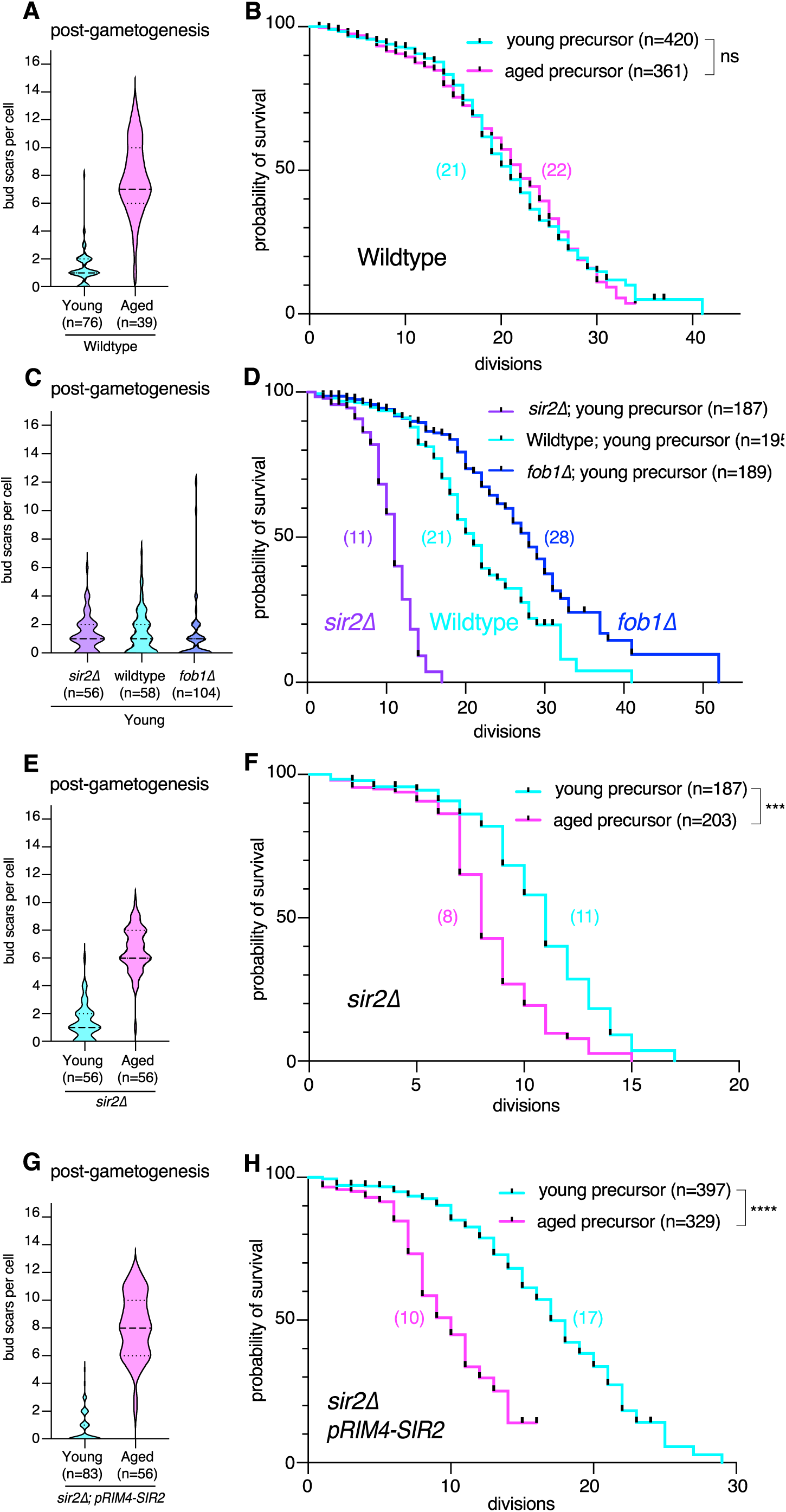
Replicative lifespan resets in wildtype gametes but rejuvenation fails in *sir2Δ* mutants through a gametogenesis-independent mechanism. (**A**) Quantification of replicative age by assessing number of bud scars in young (1.3± 1.2, mean ± SD, n=76) and aged (7.8± 2.6, mean ± SD, n=39) wildtype gamete populations. (**B**) Quantification of replicative lifespan of young (median lifespan=21, n=420, 223 censored subjects) and aged (median lifespan=22, n=361, 213 censored subjects, p=0.982 compared to young) wildtype gametes. (**C**) Quantification of replicative age by assessing number of bud scars in young *sir2Δ* (1.3± 1.3, mean ± SD, n=56), wildtype (1.4± 1.5, mean ± SD, n=104) and *fob1Δ* (11.2± 2.2, mean ± SD, n=53) gamete populations. (**D**) Quantification of replicative lifespan of young *sir2Δ* (median lifespan=11, n=187, 80 censored subjects, p<0.0001 compared to wildtype), wildtype (median lifespan=21, n=195, 103 censored subjects) and *fob1Δ* (median lifespan=28, n=286, 174 censored subjects, p<0.0001 compared to wildtype) gametes. (**E**) Quantification of replicative age by assessing number of bud scars in young (1.3± 1.3, mean ± SD, n=56, same data as shown in Fig. 4D) and aged (6.6± 1.6, mean ± SD, n=56) *sir2Δ* gamete populations. (**F**) Quantification of replicative lifespan of young (median lifespan=11, n=187, 80 censored subjects, same data as shown in Fig. 3F) and aged *sir2Δ* (median lifespan=8, n=203, 103 censored subjects, p<0.0001 compared to young) gametes. (**G**) Quantification of replicative age by assessing number of bud scars in young (0.72± 1.1, mean ± SD, n=83) and aged 8.1 ± 2.3 (mean ± SD, n=56) *pRIM4-SIR2 sir2Δ* gamete populations. (**H**) Quantification of replicative lifespan of young (median lifespan17, n=397, 266 censored subjects) and aged *pRIM4-SIR2 sir2Δ* (median lifespan=10 (n=329, 219 censored subjects, p<0.0001 compared to young) gametes. Confidence intervals were calculated using the Wilson score method and statistical significance for differences between replicative lifespans was calculated using the Log-rank (Mantel-cox) test, tick marks on survival curves represent censored events.

### Microfluidics-based analyses of gamete lifespans capture a wide range of outcomes among known short- and long-lived mutants

To test the sensitivity of our system in detecting differences in gamete replicative lifespan, we quantified lifespans of young gametes derived from known short and long-lived mutants. Deletion of the gene encoding Sir2, a NAD^+^- dependent histone deacetylase, is known to decrease replicative lifespan of yeast by approximately 50%, in part due to loss of rDNA locus silencing, leading to the generation and accumulation of extrachromosomal rDNA circles (Kaeberlein et al., 1999; Sinclair and Guarente, 1997). In contrast, deletion of the gene encoding Fob1, an rDNA spacer replication fork barrier binding protein, is reported to extend lifespan in yeast by up to approximately 40%, due to decreased rDNA recombination and thus fewer extrachromosomal rDNA circles (Defossez et al., 1999).

We sorted *sir2Δ*, wild-type, and *fob1Δ* diploid cells (**Fig. 2C**), and verified the sorting efficiency and replicative age of the young pre-gametogenic cell populations (**Fig. S3K&L**). We induced gametogenesis in the separated cell populations and found after 24 hours, 87.3% of *sir2Δ* cells, 90.4% of wildtype cells and 73.7% of *fob1Δ* cells successfully completed gametogenesis (**Fig. S3M**). Upon confirming purity of sorting and replicative age in tetrads (**Fig. 4C&S3N**), we separated the tetrads into single gametes and acquired their replicative lifespan over 72 hours. The median replicative lifespan for gametes derived from young precursor *sir2Δ* cells was 11 divisions, significantly lower than gametes derived from young wild-type cells, with a median lifespan of 21 (**Fig. 4D**, p<0.0001, Log-rank (Mantel-cox)). Gametes derived from young *fob1Δ* precursor cells, on the other hand, had a significantly increased median lifespan of 28 compared to wildtype (**Fig. 4D**, p<0.0001, Log-rank (Mantel-cox)). Thus, our microfluidic platform and workflow can capture a range of gamete lifespans from young precursor cells with mutations in established longevity-linked genes.

### Gametes lacking *SIR2* fail to reset replicative lifespan through a mechanism that is independent of gametogenesis

While testing our pipeline, we observed that gametes derived from aged *sir2Δ* precursor cells did not display complete resetting of replicative lifespan. We sorted *sir2Δ* pre-gametogenic diploid cells into a young cell population (same data as **Fig. S3K&L**) and an aged cell population (**Fig. S4A&B**). 87.3% (same data as **Fig. S3M**) of the young population and 68.0% of old the population completed gametogenesis after 24 hours (**Fig. S4C**). We verified purity and replicative age of post-gametogenic cell populations before measuring their lifespan (**Fig. 4E&S4D,** same data as shown in **Fig. S3N&4C** for young). Strikingly, the replicative lifespan of gametes derived from aged *sir2Δ* precursor cells revealed a median replicative lifespan of 8 divisions, significantly lower than for gametes derived from young *sir2Δ* precursor cells, which had a median lifespan of 11 (same data as shown in **Fig. 4D**) (**Fig. 4F**, p<0.0001, Log-rank (Mantel-cox)).

To test whether ectopic expression of *SIR2* during gametogenesis can rescue the lack of replicative lifespan resetting observed in *sir2Δ* gametes, we expressed *SIR2* ORF transgene under the control of a *RIM4* promoter fragment in *sir2Δ* cells, which resulted in gametogenesis-specific expression of Sir2 (**Fig. S4E&S4F**) (Cheng et al., 2018; Deng and Saunders, 2001). We then sorted pre-gametogenic diploid *sir2Δ* cells carrying this rescue construct into a young and aged cell population and determined completion of gametogenesis after 24h (**Fig. S4G-I**). We verified purity and replicative age of sorted cell populations upon gametogenesis before analyzing replicative lifespan of gametes (**Fig. 4G&S4J**). Gametes derived from aged *pRIM4-SIR2 sir2Δ* precursor cells showed a median replicative lifespan of 10, which was still significantly lower than gametes derived from young *pRIM4-SIR2 sir2Δ* cells, with a median lifespan of 17 (**Fig. 4H**, p<0.0001, Log-rank (Mantel-cox)). We note a slightly increased replicative lifespan of gametes derived from both young and old *pRIM4-SIR2 sir2* precursor cells compared to those derived from *sir2Δ* cells, which may be due to residual Sir2 protein observed in late gametes that could carry over through the first cell divisions (**Fig. S4F**, right most band). In summary, we show that *sir2Δ* gametes do not completely reset their lifespan following gametogenesis, however, this phenotype cannot be rescued by providing Sir2 during gametogenesis. This suggests that aged *sir2Δ* cells might accumulate defects that cannot be rescued by the gametogenic rejuvenation program.

### Selective Autophagy is necessary to reset replicative lifespan during gametogenesis

After validating our technique, we sought to determine whether quality control mechanisms known to be active during gametogenesis also promote rejuvenation. Autophagy mediates lysosomal degradation of damaged organelles and proteins (Mizushima et al., 2008). Autophagy pathways decline during aging, leading to loss of proteostasis and accumulation of cellular damage. Artificially increasing autophagy factors can lead to lifespan extension (Hansen et al., 2018; Rubinsztein et al., 2011; Simonsen et al., 2008; Tyler and Johnson, 2018). Both Atg1, an essential kinase required for initiation of autophagosome formation, and Atg8, which is required for formation of the autophagosome membrane, are upregulated during gametogenesis (**Fig. 5A&B; S5A-D**) (Cheng et al., 2018; Ichimura et al., 2000; Matsuura et al., 1997). In yeast, the general autophagy machinery is essential for meiotic entry as well as progression, which hinders evaluating the effect of bulk autophagy on gamete rejuvenation (Neiman, 2005; Tsukada and Ohsumi, 1993; Wen et al., 2016). However, Atg11, the main selective autophagy scaffold protein required for Atg1 recruitment to selective autophagy receptors on specific cargos, is also dynamically upregulated during gametogenesis (**Fig. 5C&D; S5E&F**) and we found that cells deleted for Atg11 can undergo gametogenesis (Cheng et al., 2018; Farré and Subramani, 2016; Kamber et al., 2015). Therefore, we tested whether cellular quality control via selective autophagy plays a direct role in gametogenic rejuvenation by deleting *ATG11* and measuring the rejuvenation potential with our established pipeline.

**Figure 5.**
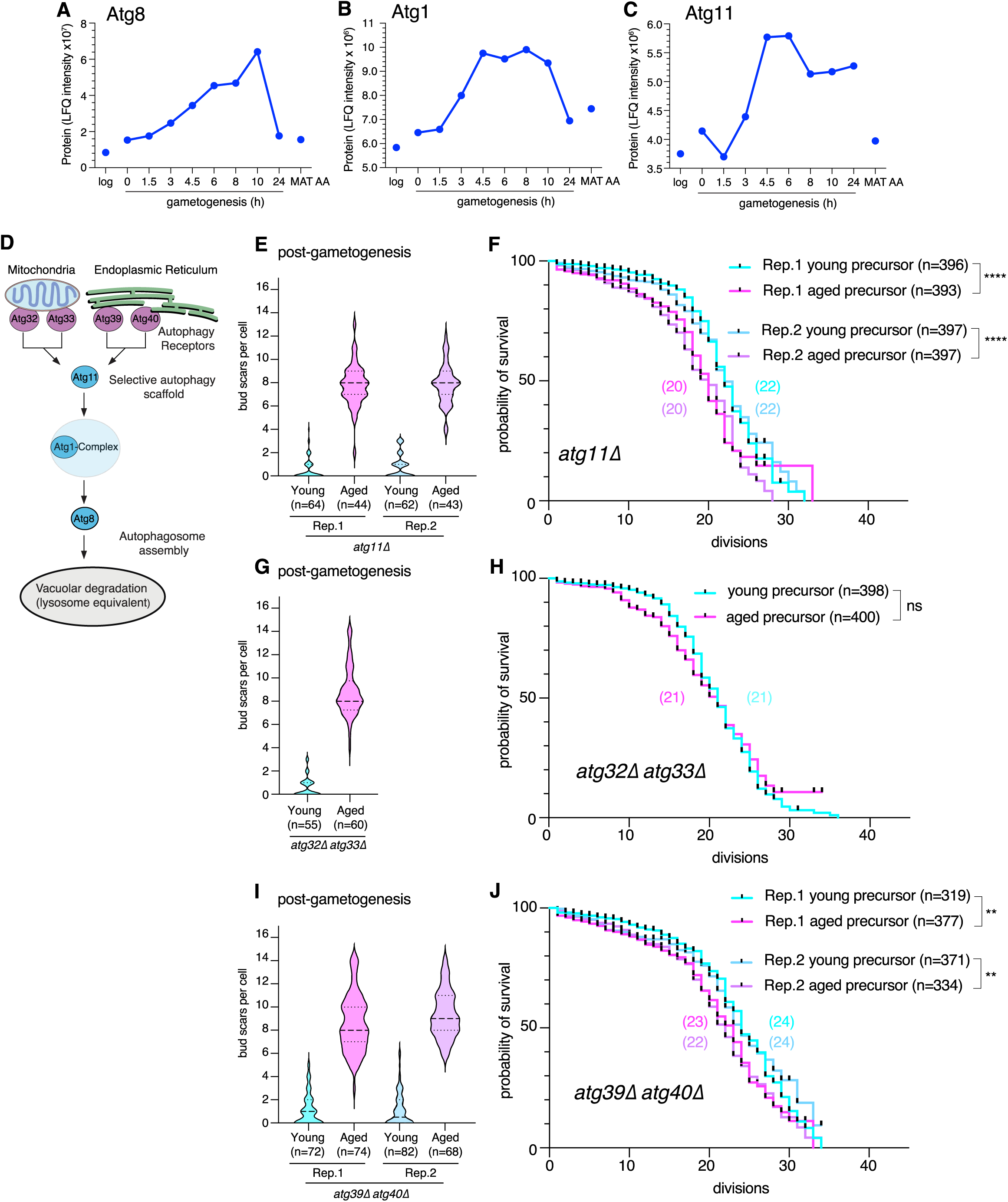
Selective autophagy of the ER but not the mitochondria is necessary to reset replicative lifespan during gametogenesis. (**A-C**) Protein levels (mass spectrometry) of autophagy factors either during log-phase vegetative growth in 2% glucose containing media (log), the indicated time in gametogenesis inducing media (h), or in control diploids that cannot undergo gametogenesis (*MAT***a**/**a**) plotted from previously generated meiotic datasets (Cheng et al., 2018). (**A**) Atg8 protein levels (mass spectrometry). (**B**) Atg1 protein levels (mass spectrometry). (**C**) Atg11 protein levels (mass spectrometry). (**D**) Schematic depiction of selective autophagy mediated by the scaffold protein Atg11 binding selective autophagy factors on specific mitochondria or ER compartments targeted for degradation, followed by autophagosome assembly and delivery to the vacuole for degradation. (**E**) Quantification of replicative age by assessing number of bud scars in *atg11Δ* tetrad populations: Rep. 1 young (0.53± 0.31, mean ± SD, n=64) and aged (7.8± 1.8, mean ± SD, n=44); Rep. 2 young (0.82± 1, mean ± SD, n=62) and aged (7.9± 1.7, mean ± SD, n=43). (**F**) Quantification of replicative lifespan of *atg11Δ* gametes: Rep.1 young (median lifespan=22, n=393, 267 censored subjects) and aged (median lifespan=20, n=396, 255 censored subjects, p>0.0001); Rep. 2 young (median lifespan=20, n=397, 280 censored subjects) and aged (median lifespan=22, n=397, 259 censored subjects, p>0.0001compared to young). (**G**) Quantification of replicative age by assessing number of bud scars in young (0.6± 0.85, mean ± SD, n=55), and aged (8.8± 2.1, mean ± SD, n=60) *atg32Δ atg33Δ* gamete populations. (**H**) Quantification of replicative lifespan of young (median lifespan=21, n=398, 154 censored subjects) and aged *atg32Δ atg33Δ* double mutants (median lifespan=21, n=400, 192 censored subjects, p=0.7414 compared to young) gametes. (**I**) Quantification of replicative age by assessing number of bud scars in *atg39Δ atg40Δ* tetrad populations: Rep. 1 young (1.1± 1.4, mean ± SD, n=72) and aged (8.6± 2.5, mean ± SD, n=74); Rep. 2 young (1.1± 1.5, mean ± SD, n=82) and aged (9.4± 2, mean ± SD, n=68). (**J**) Quantification of replicative lifespan of *atg39Δ atg40Δ* gametes: Rep.1 young (median lifespan=24, n=319, 195 censored subjects) and aged (median lifespan=23, n=377, 238 censored subjects, p=0.0030 compared to young); Rep. 2 young (median lifespan=24, n=371, 240 censored subjects) and aged (median lifespan=22, n=334, 219 censored subjects, p=0.0080 compared to young). Statistical significance for differences between replicative lifespans was calculated using the Log-rank (Mantel-cox) test, tick marks on survival curves represent censored events.

We sorted pre-gametogenic cells into young and aged cell populations and 24h after induction verified the efficiency of gametogenesis (**Fig. S5G-I**). Replicative age of tetrad population was confirmed by quantifying bud scars per tetrad and number of biotinylated tetrads in the population (**Fig. 5E&S5J**). Strikingly, when quantifying the replicative lifespan of gametes, we found a significant difference between gametes from aged and young *atg11Δ* precursor cells that was highly reproducible (**Fig. 5F**, Rep.1, p<0.0001, Rep.2, p<0.0001, Log-rank (Mantel-cox), compared to **Fig. 4B**). This result suggests that Atg11-mediated selective autophagy is essential for full rejuvenation of gametes derived from aged precursor cells.

### Mitophagy is not necessary to reset replicative lifespan during gametogenesis

Atg11 recruits the autophagy machinery upon binding to specific selective autophagy receptors that can mark select organelles for degradation (**Fig. 5D**), thus we wondered if selective autophagy of a specific compartment is necessary for gametogenic rejuvenation. To address this, we first investigated a potential role for Atg32, an autophagy receptor localized on the outer mitochondria membrane that mediates selective removal of damaged and excess mitochondria via mitophagy (Kanki et al., 2009; Kondo-Okamoto et al., 2012; Okamoto et al., 2009). Atg32 is strongly upregulated in mid-meiosis (**Fig. S6A-C**) (Cheng et al., 2018) and specific mitochondrial remodeling and inheritance during gametogenesis has been described before (Sawyer et al., 2018; Suda et al., 2007).

We deleted *ATG32* in diploid cells and tested their gametogenic rejuvenation potential. We again sorted pre-gametogenic cells into young and aged cell populations, determined efficiency of gametogenesis and analyzed biotin label and replicative age of post gametogenic cell populations (**Fig. S6D-H**). No significant difference of replicative lifespan was observed between gametes from aged compared to young *atg32Δ* precursor populations (**Fig. S6I**, p=0.2691, Log-rank (Mantel-cox)). Atg33, which is localized on the outer mitochondria membrane also confers selectivity during mitophagy under certain growth conditions and is upregulated during meiosis (**Fig. S6J-L**). To exclude the possibility that *ATG33-*mediated mitophagy acts in parallel thereby masking a possible rejuvenation phenotype of *atg32Δ* mutants, we constructed *atg32Δ atg33Δ* double mutants and tested their rejuvenation potential. We age-sorted *atg32Δ atg33Δ* pre-gametogenic cells into young and aged cell populations, verified efficiency of gametogenesis and analyzed biotin labelling and replicative age of post gametogenic cell populations (**Fig. 5G&S6M-P**). As for *atg32Δ* single mutants, no significant difference in replicative lifespan was observed in *atg32Δ atg33Δ* gametes (**Fig. 5H**, p=0.7414, Log-rank (Mantel-cox)). These experiments suggest that selective mitophagy is dispensable for gametogenic rejuvenation. Failure of *atg32Δ atg33Δ* to phenocopy *atg11Δ* implies that a different specific arm of selective autophagy mediated by Atg11 is necessary for gametogenic rejuvenation.

### ER-phagy is necessary for full gametogenic lifespan resetting

It has previously been shown that Atg11 dependent selective autophagy mediated by the autophagy receptors Atg40, and to a lesser extent Atg39, control ER inheritance during gametogenesis. Atg39 and Atg40 are upregulated during gametogenesis (**Fig. S7A-F**) (Cheng et al., 2018) and meiotic remodeling of the ER combined with ER-phagy prevent specific subpopulations of the ER from being passed onto gametes. It has been proposed that these pathways constitute a quality control mechanism to ensure gametogenic rejuvenation, however this has not been directly tested (Otto et al., 2021; Otto and Brar, 2022).

To determine if Atg39, a transmembrane autophagy receptor in the perinuclear ER mediating autophagy, impacts gametogenic rejuvenation, we first sorted diploid *atg39Δ* cells into a young and aged cell population (**Fig. S7G&H**) (Mochida et al., 2015). In the young and aged populations, 70.0% and 82.0% completed gametogenesis, respectively (**Fig S7I**). We verified age separation of post-gametogenic populations before measuring lifespan of gametes with our microfluidic devices (**Fig. S7J&K**). We found no significant difference for replicative lifespan of *atg39Δ* gametes derived from aged or young precursors (**Fig. S7L**, p=0.418, Log-rank (Mantel-cox)).

Next, we checked if gametogenic reset of replicative lifespan is impaired in cells lacking Atg40, an autophagy receptor localized at the cortical and cytoplasmic ER mediating fragmentation and encapsulation of ER subdomains (Mochida et al., 2015). Pre-gametogenic *atg40Δ* mutant cells were sorted into young and aged cell populations, efficiency of gametogenesis as well as biotin label and replicative age of post-gametogenic cell populations was determined upon gametogenesis (**Fig S7M-Q**). Similar to *atg39Δ* cells, no significant difference in replicative lifespan was observed between *atg40Δ* gametes derived from aged or young precursors (**Fig. S7R**, p=0.389, Log-rank (Mantel-cox)).

Because there is known functional redundancy between Atg39 and Atg40 (Mochida et al., 2015; Otto et al., 2021), we tested if *atg39Δ atg40Δ* cells were still capable of completely resetting replicative lifespan during gametogenesis. We verified purity and replicative age upon sorting *atg39Δ atg40Δ* diploid cells into young and aged populations (**Fig. S8A&B**, Rep.1). Upon gametogenesis, we determined efficiency of gametogenesis and assured sorting purity as well as replicative age (**Fig 5I; S8C&D**, Rep.1). Excitingly, lifespan analysis of aged gametes from aged *atg39Δ atg40Δ* precursor cells revealed a significantly reduced median replicative lifespan compared to gametes from young precursor cells (**Fig 5J**, Rep.1, p=0.0030, Log-rank (Mantel-cox)). The failure to fully rejuvenate in aged *atg39Δ atg40Δ* gametes was highly reproducible (**Fig. 5I&J**, Rep.2, p=0.0080, Log-rank (Mantel-cox); **S8A-D**). Together, these data revealed that Atg39 and Atg40 both contribute through partially redundant mechanisms to gametogenic rejuvenation, and that full replicative lifespan resetting during gametogenesis depends on the presence of these selective ER-phagy receptors.

### Meiosis specific expression of *ATG40* is sufficient to rescue resetting of replicative lifespan in aged ER-phagy-deficient cells

To determine whether this difference in replicative lifespan between gametes from aged and young *atg39Δ atg40Δ* precursor cells is due to lack of selective ER-phagy specifically during gametogenesis, we tested if ectopic meiotic expression of *ATG40* is sufficient to rescue it. We constructed a *atg39Δ atg40Δ* double mutant with the *ATG40* ORF under the control of a *NDT80* promoter fragment, which displays meiosis-specific expression (**Fig. S9A**) as well as a *atg39Δ atg40Δ* double mutant control strain (Xu et al., 1995). We confirmed gametogenesis-specific expression by immunoblotting (**Fig. S9B**) and performed rejuvenation assays in independent duplicates.

First, we sorted pre-gametogenic *atg39Δ atg40Δ* control cells into young and aged cell populations and determined efficiency of gametogenesis (**Fig. S9C-E**). We then analyzed the percentage of biotin labeled tetrads and replicative age of post-gametogenic cell populations (**Fig 6A, S9F**). Lifespan analysis confirmed again a similar significant and reproducible lower median replicative lifespan in gametes from aged precursor *atg39Δ atg40Δ* control cells compared to gametes from young precursor cells (**Fig. 6B**, Rep. 1, p<0.0001, Rep. 2, p<0.0001, Log-rank (Mantel-cox)). Next, we sorted pre-gametogenic *atg39Δ atg40Δ pNDT80-ATG40* cells into young and aged cell populations, determined efficiency of gametogenesis and confirmed construct expression (**Fig. S8G-K**). After completion of gametogenesis, we confirmed the percentage of biotinylated tetrads and mean replicative age of young and aged populations (**Fig 6C&S9L**). Excitingly, there was reproducibly no significant difference of replicative lifespan in gametes from aged precursor *atg39Δ atg40Δ pNDT80-ATG40* cells, (**Fig. 6D**, Rep.1, p=0.4519, Rep.2, p=0.9538, Log-rank (Mantel-cox)). Finally, we quantified ER dynamics during gametogenesis in young and aged wildtype and *atg39Δ atg40Δ* mutant cells (**Fig. S10A&B**) by probing for endogenously-tagged Sec63-GFP, an abundant ER membrane protein present in both ER sheets and tubules. We found that Sec63-GFP accumulates during gametogenesis in young and aged cells lacking *atg39Δ atg40Δ,* relative to wildtype cells, indicating that ER turnover through ER-phagy significantly impacts the amount of ER inherited by gametes (**Fig. 6E&F, S10C-F).** Together, these experiments demonstrate that complete resetting of replicative lifespan during gametogenesis is dependent on selective ER-phagy, possibly due to programmed turnover of specific ER species during gametogenesis (**Fig. 6G**).

**Figure 6.**
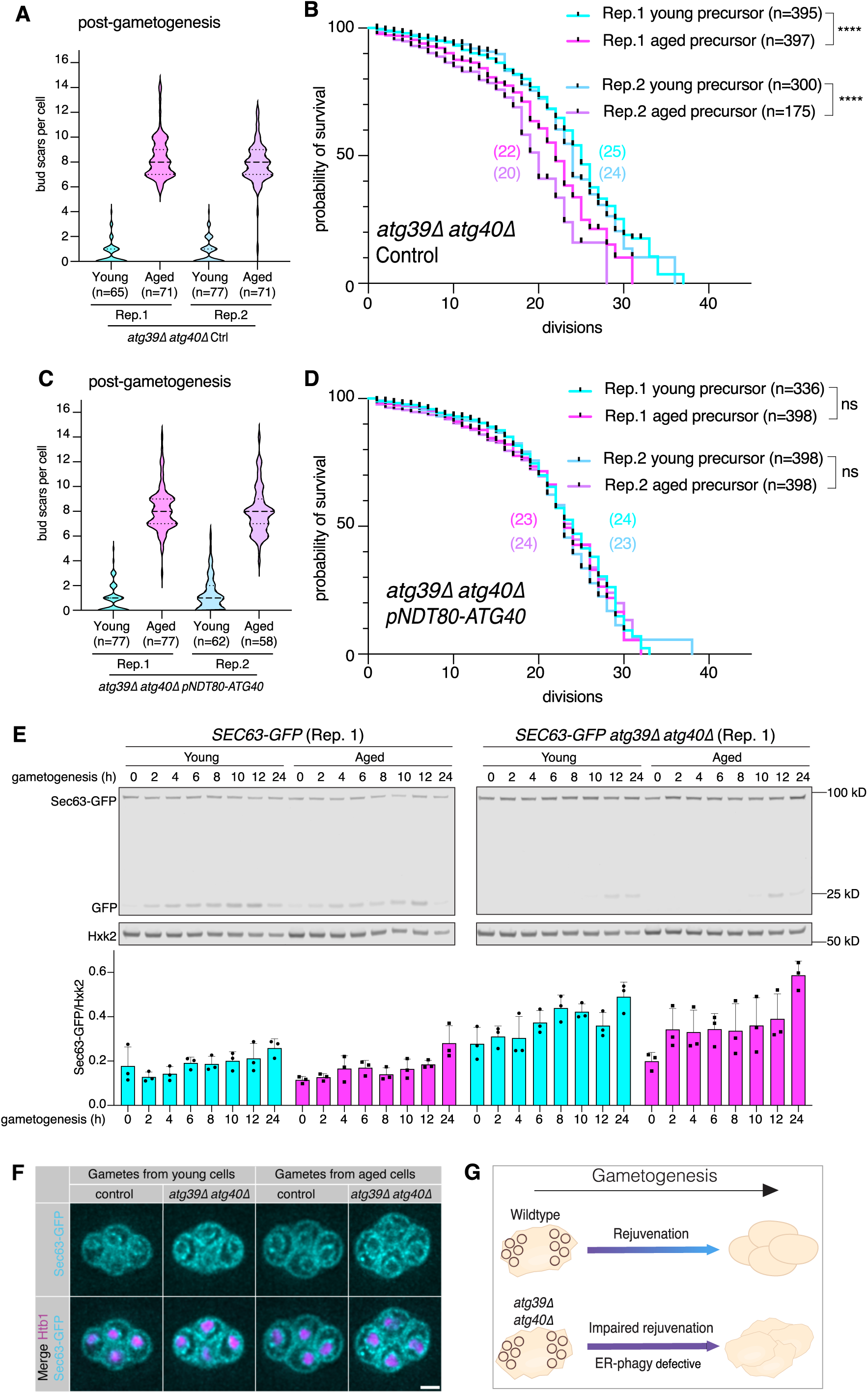
Expression of *ATG40* during gametogenesis is sufficient to rescue replicative lifespan resetting in cells that lack ER-phagy receptors. (**A**) Quantification of replicative age by assessing number of bud scars in *atg39Δ atg40Δ* control tetrad populations: Rep. 1 young (0.62± 0.92, mean ± SD, n=65) and aged (8.4± 1.7, mean ± SD, n=71); Rep. 2 young (0.74± 0.97, mean ± SD, n=77) and aged (7.8± 1.6, mean ± SD, n=71). (**B**) Quantification of replicative lifespan of *atg39Δ atg40Δ* control gametes: Rep.1 young (median lifespan=25, n=397, 229 censored subjects,) and aged (median lifespan=22, n=395, 277 censored subjects, p<0.0001 compared to young); Rep. 2 young (median lifespan=24, n=300, 185 censored subjects, 1 technical replicate) and aged (median lifespan=20, n=175, 122 censored subjects, 1 technical replicate, p<0.0001 compared to young). (**C**) Quantification of replicative age by assessing number of bud scars in *atg39Δ atg40Δ pNDT80-ATG40* tetrad populations: Rep. 1 young (0.87± 1.1, mean ± SD, n=77) and aged (8.1± 1.8, mean ± SD, n=77); Rep. 2 young (1.2± 1.47, mean ± SD, n=62) and aged (8.0± 1.9, mean ± SD, n=58). (**D**) Quantification of replicative lifespan of *atg39Δ atg40Δ pNDT80-ATG40* gametes: Rep.1 young (median lifespan=24, n=398, 231 censored subjects) and aged (median lifespan=23, n=336, 242 censored subjects, one technical replica, p<0.4519 compared to young); Rep. 2 young (median lifespan=23, n=398, 256 censored subjects) and aged (median lifespan=24, n=398, 281 censored subjects, p=0.9538 compared to young). **E**) Immunoblot of protein samples form young and aged *SEC63-GFP* (left) and *atg39Δ atg40Δ SEC63-GFP* (right) cells to quantify protein levels (anti-GFP) and loading control (Hxk2) at different timepoints upon transfer to gametogenesis-inducing media (gametogenesis (h)) and quantification of Sec63-GFP levels normalize to loading control (Hxk2), bar graphs show mean error bars show ± SD, 3 independent replicas. (**F**) Representative single z-slices of post-gametogenic young (left panel) and aged (right panel) *SEC63-GFP* and *atg39Δ atg40Δ SEC63-GFP* tetrads respectively, visualizing Sec63-GFP (cyan) and Htb1-mCherry (magenta). Scale bar is 2 µm. (**G**) Schematic depiction of impaired gametogenic rejuvenation in ER-phagy mutants. Statistical significance for differences between replicative lifespans was calculated using the Log-rank (Mantel-cox) test, tick marks on survival curves represent censored events.

## Discussion

Here, we present a high throughput microfluidics-based platform to identify the molecular mechanisms underlying the natural cellular rejuvenation that accompanies gametogenesis in budding yeast cells. This technique accelerates investigations of gametogenic rejuvenation efficiency and will be instrumental in revealing further mechanistic insights into the conserved process of gametogenic rejuvenation. Using our method, we corroborate previous reports using microdissection that replicative lifespan is completely resets by gametogenesis (Ünal et al., 2011). Analysis of gamete lifespan in known short- and long-lived mutants reveals a large dynamic range of measurement using this approach. We found that in *sir2Δ* gametes from aged precursor cells, replicative lifespan was not fully reset due to a gametogenesis-independent mechanism. Finally, using this method we revealed that while mitophagy appears to be dispensable for gametogenic rejuvenation, Atg39- and Atg40-dependent ER-phagy is necessary for gametogenic rejuvenation. The reconstitution of Atg40 during meiosis in this background is sufficient to completely reset replicative lifespan of aged gametes.

The custom microfluidic devices presented here, with an open U-trap design, can capture a wide dynamic range of replicative lifespan with lower confinement of cells in space compared to other trap designs that can lead to negative effects on longevity due to compression of cells (Gao et al., 2020). This open design does present the drawback that cells might undergo ejection prematurely before senescence can be observed, due to either a significant increase of cell size during the last few divisions of replicative life, or due to minor pressure aberrations in the fluidics system over time. To avoid biases when evaluating replicative lifespan of gametes, we counted every gamete that was loaded at the start of our time lapse and classified cells that ejected as censored subjects during analysis (see Methods). Following this approach, we can consistently and robustly measure replicative lifespan. We find the quantified increase of 33.33% in median lifespan of young *fob1Δ* gametes and 52% decrease for *sir2Δ* compared to lifespan of young wild-type gametes to be within the range of previous reports (Kaeberlein et al., 2004; Defossez et al., 1999; Kaeberlein et al., 1999).

*SIR2* is a well-characterized conserved histone deacetylase of the Sirtuin family and regulates replicative lifespan by silencing the ribosomal DNA locus (Kaeberlein et al., 1999; Sinclair and Guarente, 1997). Deletion of the *SIR2* leads to increased rDNA recombination and broad chromosome instabilities due to destabilization of the rDNA locus (Ide et al., 2010; Kobayashi et al., 2004; Kobayashi and Ganley, 2005). Gametes from aged *sir2Δ* precursor cells failed to reset replicative lifespan, a phenotype that could not be rescued by gametogenesis-specific expression of Sir2. These results indicate that Sir2 does not play a direct role during gametogenic rejuvenation itself. One plausible explanation for this is that due to elevated recombination, aged *sir2Δ* cells accumulate chromosomal aberrations, which cannot be corrected during gametogenesis, and are propagated to gametes, thus reducing their life expectancy. Elevated pre-existing damage in *sir2Δ* cells is consistent with the substantially reduced efficiency of gametogenesis in aged cells in this background (**Fig. S4C**) (Boselli et al., 2009), which was not generally observed in other backgrounds we analyzed.

Autophagy of specific cellular components is known to be an important mechanism for cells to counteract age-associated damage and delay senescence (Hansen et al., 2018; Rubinsztein et al., 2011; Simonsen et al., 2008; Tyler and Johnson, 2018). The general autophagy machinery is upregulated during gametogenesis and is necessary for meiotic entry and progression (Neiman, 2005; Wen et al., 2016). We found that selective autophagy is a crucial pathway to mediate gametogenic rejuvenation and show specificity of this process as ER-phagy but not mitophagy is particularly important for gamete rejuvenation.

The mitochondrial network undergoes substantial remodeling during gametogenesis (Sawyer et al., 2018; Suda et al., 2007). However, gametes from aged *atg32Δ atg33Δ* precursor cells completely reset replicative lifespan, indicating that Atg32- and Atg33-mediated selective mitophagy is not necessary for gametogenic rejuvenation. These findings suggest that dedicated mitochondrial quality control via selective autophagy may be dispensable for gametogenic rejuvenation. Autophagy-independent mechanisms, such as potential sequestration of damaged mitochondria via the gametogenic uninherited compartment may ensure inheritance of healthy mitochondria (King et al., 2019; Ruediger et al., 2025). This will be interesting to address in future studies using our method.

Developmentally programmed ER remodeling followed by selective autophagy of ER components regulates ER inheritance during yeast gametogenesis (Otto et al., 2021; Otto and Brar, 2022). ER-specific autophagy receptors Atg40 and Atg39, localized on the cortical ER and perinuclear ER, respectively, control this process (Mochida et al., 2015). *atg40Δ* but not *atg39Δ* cells fail to degrade cortical ER components via autophagy during gametogenesis. However, *atg39Δ* cells show reduced autophagy-mediated loss of cortical and perinuclear ER proteins, suggesting functional redundancy with Atg40. Intriguingly, *atg40Δ atg39Δ* double mutants have an additive effect on ER protein degradation (Otto et al., 2021). We show that aged *atg40Δ atg39Δ* precursor cells accumulate ER components during gametogenesis and fail to produce gametes with a fully reset lifespan. Loss of Rer1, a membrane protein retrieving receptor that maintains ER compartmentalization, has been shown to extend replicative lifespan in vegetative yeast through elevated autophagy triggered by ER stress (Ghavidel et al., 2015; Sato et al., 1997). We speculate that the observed rejuvenating defect of ER-phagy-deficient cells could be caused by a failure of meiotic clearance of damaged and aggregated ER components that accumulated in aged cells (Mochida et al., 2015; Wilkinson, 2020), an interesting area to investigate in future studies.

ER-phagy is functionally conserved from yeast to metazoans and essential to maintain ER quality (Dikic and Elazar, 2018). In mammals, mutations in ER-phagy receptor FAM134B cause hereditary sensory neuropathy and impaired ER turnover has been linked to neurodegeneration, including Parkinson‗s disease (Kim et al., 2023; Murphy et al., 2012). Thus, ER quality control mediated by autophagy is a conserved mechanism to preserve cellular fitness. Our finding suggests that meiotic ER-phagy contributes to gamete rejuvenation by assuring ER integrity.

The technique presented here significantly increases throughput and makes it feasible to identify molecular factors that mediate cellular rejuvenation during the conserved process of gametogenesis. To gain more mechanistic and temporal understanding, this assay can easily be combined with auxin-inducible degradation of factors of interest and simultaneous imaging of established aging biomarkers in gametes that fail to completely rejuvenate. An important future goal will be to determine if newly identified rejuvenation factors that control gamete quality are sufficient to extend replicative lifespan in mitotic cells, and to test if they have a conserved function in combating aging in other systems.

## Materials and Methods

### Yeast strains and plasmids

Standard molecular methods were used to transform and generate all yeast strains, which were derived from the SK1 background. Specific genotypes, plasmids and primers used to generate strains are listed in Supplementary information (**Supplementary tables 1-3**). Deletions were generated by homology directed replacement of ORFs with selection cassettes (Longtine et al., 1998; Powers et al., 2022). The *pPMA2-PMA2-GFP* transgene plasmid was generated by Gibson assembly of the *PMA2* promoter and ORF region, amplified from genomic DNA, with an in frame *GFP*-containing *TRP1* single integration vector (Gibson et al., 2009). The *pTetO7.1-STE5* promoter swap was generated by amplifying a *NatR- pTetO7.1* fragment using primers that introduced homology for the *STE5* locus (Bellí et al., 1998; Longtine et al., 1998). The *pRIM4-SIR2-V5* rescue construct was generated by Gibson assembly of the *SIR2* ORF region with *pRIM4* and *V5* containing *LEU2* single integration vector (Gibson et al., 2009). The *pNDT80-ATG40-V5* rescue construct was generated by Gibson assembly of the *ATG40* ORF region with *pNDT80* and *V5* containing *LEU2* single integration vector (Gibson et al., 2009). Generated plasmids were assembled using HiFi DNA assembly mix (NEbuilder HiFi DNA Assembly Master Mix, E2621L). *pRNR2-TetR-TUP1, PTetO7.1-TetR* adapted from (Azizoglu et al., 2021) was cloned into a *URA3* single integration vector by Gibson assembly (Gibson et al., 2009). Single integration plasmids were linearized with PmeI (New England Biolabs) mediated restriction digest before transformations and integrations were verified by PCR and sanger sequencing. Generated strains and plasmids are available upon request. The following alleles were constructed in previous studies: *atg39Δ, atg40Δ, ATG40-3V5, SEC63-eGFP* (Otto et al., 2021)*, HTB1-mCherry (Matos et al., 2008), flo8Δ, amn1^D368V^(Tina L Sing et al., 2022)*.

### Media and growth conditions

Unless otherwise indicated, diploid cells were grown in YPD (1% yeast extract, 2% peptone, 2% glucose, 22.4 mg/L uracil, and 80 mg/L tryptophan). To induce gametogenesis cells were transferred to conditioned sporulation media containing 2% potassium acetate, amino acid supplements and ampicillin (40 mg/L adenine, 40 mg/L uracil, 10 mg/L histidine, 10 mg/L leucine and 10 mg/L tryptophan, 100 μg/mL ampicillin) at an OD_600_ of 0.8. All meiotic cultures were incubated on a shaker for 24h at 30°C to complete gametogenesis. To increase efficiency of gametogenesis, conditioned sporulation media was obtained by filter sterilizing a meiotic culture at 1.85 OD_600_ after 5 hours at 30°C and was stored at 4°C. For immunoblotting saturated YPD 2% cultures were re-diluted to 0.2 OD_600_ in BYTA (1% yeast extract, 2% bacto tryptone, 1% potassium acetate, and 50mM potassium phthalate) containing acetate as a carbon source and then grown overnight to saturation at 30°C before transfer to unconditioned sporulation media without ampicillin at an OD_600_ of 1.85.

### Fluorescent Microscopy

Bud scar staining and Pma2-GFP expression patterns during meiosis and gamete maturation were visualized using a DeltaVision Elite wide-field fluorescence microscope (GE Healthcare), equipped with a 60x/1.42 NA oil-immersion objective. Images were deconvolved with softWoRx imaging software (GE Healthcare). Time lapse images were acquired using a CellAsics system as previously described (King et al., 2019). Replicative lifespan timelapse imaging was performed using an ECHO Revolution (RON-K) microscope equipped with an Olympus 40x/1.4 NA oil-immersion objective.

### Age sorting of pre-gametogenic yeast cells

Young and aged cell populations were sorted by biotin-labeling of a founder population followed by bead-mediated magnetic sorting (Smeal et al., 1996). First, a patch of diploid cells was inoculated in YPD (2%) and grown overnight shaking at room temperature until saturation. Cells were then re-diluted in YPD (2%) to an OD_600_ of 0.2 and grown at 30°C to an OD_600_ of 0.6-1. To increase yield, 2 x 8 OD_600_ units were harvested per genotype, washed 3x with PBS (pH 8), and then were transferred into PBS (pH 8) containing 2 mg/mL EZ-Link Sulfo-NHS-LC-biotin (ThermoFisher Scientific). Cells were vortexed for one minute and incubated for 30 minutes at 4°C on a nutator, washed 4 times with PBS + glycine (pH 8 + 100 mM glycine), and then washed once with YPD 2% before being added to two separate 500 mL flasks containing 50 mL YPD 2% + ampicillin (100 μg/mL) each. Biotin labeled cultures were aged for 15-19h shaking at 30°C. Aged cell cultures were pelleted, washed once with 20 mL PBS (pH 7.4 + 0.5% BSA), and then resuspended in 20 mL PBS (pH 7.4 + 0.5%) BSA containing 100 μL of anti-biotin magnetic beads (Miltenyi Biotechnology). Cells were incubated for 15 min at 4°C and mixed regularly by inversion, washed once with 20 mL PBS (pH 7.4 + 0.5% BSA) and then resuspended in 5 mL PBS (pH 7.4 + 0.5% BSA). Using a QuadroMacs sorter, both suspensions were applied to a column each. The first flowthrough from each column was collected, combined, and used as the young cell population. Each column was washed 4 times with 3 mL PBS (pH 7.4 + 0.5% BSA) before eluting into a second column, to increase purity. The second columns were washed three times with 3 mL PBS (pH 7.4 + 0.5% BSA), cells were eluted, combined, and subsequently used as the aged cell population.

### Immunostaining and gametogenesis

Young and aged cell suspensions were washed twice with 12 mL PBS (pH 7) and resuspended in 4 mL PBS (pH 7). 4 μL of Wheat Germ Agglutinin, Alexa 350 Fluor conjugate (1 μg/mL, ThermoFisher Scientific) and anti-streptavidin (564 nm) (1 μg/mL) was added respectively and incubated for 15 minutes in the dark at room temperature. Cells where washed (3x) with 12 mL of unconditioned sporulation media and resuspended to an OD_600_ of 0.8 in conditioned sporulation media + ampicillin (100 μg/mL). 0.1 OD_600_ units of cells was fixed in 3.7% formaldehyde to assess age before gametogenesis. Cultures were incubated for 24 hours at 30°C in the dark to complete gametogenesis and a 0.1 OD_600_ units of cells were used directly to count tetrad formation and another fraction of cells were fixed in 3.7% formaldehyde to assess age post-gametogenesis by imaging and counting biotinylated cells as well as bud scars.

### Tetrad separation into single gametes and re-growth

Post-gametogenic age sorted cultures were pelleted and resuspended in 900 μL H_2_O, 100 μL Zymolyase (1 mg/mL) and 2 μL B-mercaptoethanol and the suspension was incubated for 3 hours on a nutator at 30°C. Cells were washed (1x) with H_2_O and resuspended in 1 mL H_2_O followed by repeated sonication (6x 20 seconds) incubating the cells 2-3 minutes on ice in- between sonication cycles. Upon tetrad separation, separated gametes were filtered (30 µm Pre-separation filter, militenyibiotec) and diluted in YPD 2% + ampicillin (100 μg/mL) to an OD_600_ of 0.8. After 3 hours at incubation on a nutator at 30°C, 1 mL of cell suspension was aliquoted, sonicated briefly for 5 seconds and filtered before loading into the microfluidics system.

### Microfluidics loading and timelapse imaging of replicative lifespan

Microfluidics devices (**Fig S1** adapted from (Jo et al., 2015);18++ chips from iBiochips) were setup using the manufacturer’s recommendations on an ECHO Revolution fluorescence microscope in the inverted position with the environmental chamber set to 30°C. Briefly, 20 mL syringes (Becton, Dickinson and Company) were filled with ∼15 mL of YPD 2% + ampicillin (100 μg/mL) + 1% Pluronic F-127 (P2443, Sigma), and then connected to a Clay Adams Intramedic Luer-Stub Adapter (Becton, Dickinson and Company). Non-DEHP Medical Grade Tubing (iBiochips, MTB-100, ID = 0.202”, OD = 0.060”, wall = 0.20”) equipped with a hollow stainless-steel pin (iBiochips, HSP-200) was then used to connect the media syringe to the imaging units using the “media loading” ports. “Cell loading” ports were plugged with solid stainless-steel pins (iBiochips, SSP-200) while imaging units were primed with media for at least 2 hours with a flow rate of 2.4 μL/min using micropumps (iBiochips,78-7200BIO). After all units were filled with media, pumps were stopped an hour prior to cell loading. For cell loading, 5 mL syringes (Becton, Dickinson and Company) with Clay Adams Intramedic Luer-Stub Adapter (Becton, Dickinson and Company) were filled with 1 mL of cells diluted to an OD_600_ of 0.8 in YPD 2% + ampicillin (100 μg/mL). Solid stainless-steel pins were then removed from the “cell loading” ports and tubing and open stainless-steel pins were used to connect the syringe containing cells with the microfluidic device. Cells were loaded manually and, once sufficient cells were loaded, the “cell loading” ports were plugged with solid stainless-steel pins and media flow was resumed at 2.4 μL/min.

Timelapse imaging was performed using an Olympus 40x/1.4 NA oil-immersion objective with intervals of 15-20 minutes for 72 hours. The x-, y-, and z-motors were set to medium speed; however, varying stage speeds caused sporadic differences in imaging intervals. Thus, the recorded acquisition time provides accurate timing information. For each chamber, 2x27 images and three z-slices per position with 3-5 μm spacing, to compensate for focal plane differences across a single unit, were acquired. Autofocus was set across a 20 μm range with 0.2 μm spacing using the “Course-Fine” setting (Threshold = 4, Contrast Limit = 4). The XY Stage and Z Motor were both set to medium speed and Hyperscan was enabled for the “BF” channel using “Y-Axis” Axis Priority and “Bi” Scanning Pattern.

### Quantification of replicative age and purity of sorted cells

Z-stacks (36 slices, 0.2 µm spacing) of fixed young and aged cells/tetrads were acquired on a DeltaVision Elite wide-field fluorescence microscope (GE Healthcare), using a 60x/1.42 NA oil-immersion objective and deconvolved with softWoRx imaging software (GE Healthcare). To reliably count number of bud scars, images were cropped, 3D projected and counted using a custom semi-automated FIJI script (ImageJ2, V. 2.14.0/1.54m, (Schindelin et al., 2012)). Max intensity Z-projections were generated and displayed as examples in figures.

### Quantification of gamete replicative lifespan

After the movie was completed, the ECHO batch exporter tool was used to export stitched TIFF images. Individual TIFF images were converted to 8-bit using FIJI (ImageJ2, V. 2.14.0/1.54m, (Schindelin et al., 2012)), then full imaging unit time-lapse could be opened using the “Bio Formats Import” tool with the “Group files with similar names” box checked. Single Pma2-GFP positive cells (gametes) were identified by eye and cropped both in the FITC image and the brightfield time course using a custom semi-automated FIJI script. Next using a semi-automated FIJI script, the replicative lifespan of Pma2-GFP positive cells (gametes) identified in the FITC image were counted, verifying identity by comparing the FITC and brightfield time-lapse for each single gamete. If a cell ejected from the U-trap before senescence was observed, this was noted and treated as a censored subject for quantification. For most experiments, counts from two technical replicates were concatenated unless otherwise indicated. Although throughput to quantify gametogenic reset of lifespan is significantly higher than with previous methods, cell divisions of each gamete must still be counted manually by a trained scientist rendering data analysis as a current bottleneck of the presented method.

### Immunoblotting

For each sample, 1.85-10 OD_600_ units were harvested and pelleted from either log-phase YPD 2%, saturated YPD 2%, saturated BYTA or indicated timepoints upon transfer to sporulation media. For each sample, 1-2 mL 5% trichloroacetic acid was added and cells were incubated for ≥10 minutes at 4°C minutes, washed (1x) with acetone, and pellets was dried overnight. For protein extraction, ∼100 μL glass beads and 100 μL of lysis buffer (50 mM Tris-HCl pH 8.0, 1 mM EDTA, 3 mM DTT, 2.2 mM PMSF, 1x complete EDTA-free inhibitor cocktail (Roche)) were added to the dried pellets and transferred to a Mini-Beadbeater-96 (BioSpec) for 5 minutes. Fifty μL of 3X SDS sample buffer (187.5 mM Tris pH 6.8, 6% β-mercaptoethanol, 30% glycerol, 9% SDS, 0.05% bromophenol blue) was added and samples were for boiled for 5 minutes at 95°C. Polyacrylamide gel electrophoresis (4–12% Bis-Tris Bolt gels, Thermo Fisher) and semi-dry transfer (Trans-Blot Turbo Transfer System, Bio-Rad) onto nitrocellulose membranes apparatus was used to separate and visualize proteins respectively. Odyssey PBS Blocking Buffer (LI-COR Biosciences) was used to block membranes (90 minutes, room temperature), and membranes were incubated overnight at 4°C with either a mouse anti-3v5 (R960-25; RRID: AB_2556564, Thermo Fisher) antibody at 1:2000 in PBS +0.1% Tween-20 or a mouse anti-GFP (JL8, Takara) at 1:2000 in PBS +0.1% tween and a rabbit anti-Hx2k (RRID:AB_219918, 100–4159, Rockland) at 1:20000 in PBS +0.1% tween. Membranes were washed with PBS+ 0.1% Tween-20 (4x, 5 minutes each) and incubated with secondary antibody solutions containing anti-mouse conjugated to IRDye 800CW at a 1:15,000 dilution (RRID:AB_621847, 926–32212, LI-COR Biosciences) and an anti-rabbit conjugated to IRDye 680RD at a 1:15,000 dilution (RRID:AB_10956166, 926–68071, LI-COR Biosciences) in PBS +0.1% Tween-20. Membranes were imaged using an Odyssey CLx system (LI-COR Biosciences).

### Statistical analysis

Prism Graphpad (version 10.1.0) was used to plot and analyze data. Prism Graphpad was used to calculated mean and standard deviation of replicative ages and to assess statistical significance in differences between replicative lifespans using the Log-rank (Mantel-cox) test. A Python script was used to calculated 95% confidence intervals using the Wilson score method.

## Acknowledgements

We thank the current and former members of the Brar and Ünal laboratory for critical discussion and comments on the manuscript. This work is supported by the Swiss National Science Foundation mobility fellowship P500PB_214421 to S.S., G.B. is supported by funding from the National Institutes of Health (R01AG071869) and the Astera Fund, TLS was supported by a Longevity Impetus Grant from the Norn Group, Hevolution Foundation and Rosenkranz Foundation. E.Ü. is supported by funding from the National Institutes of Health (R01AG071801) and the Astera Institute.

## Author contributions

S.S. and N.P. performed experiments, S.S. and N.P. analyzed the data, S.S., T.L.S., N.P., J.S.G., E.Ü., G.A.B., wrote and edited the manuscript.

## Supplementary Figures & Tables

**Figure S1.**
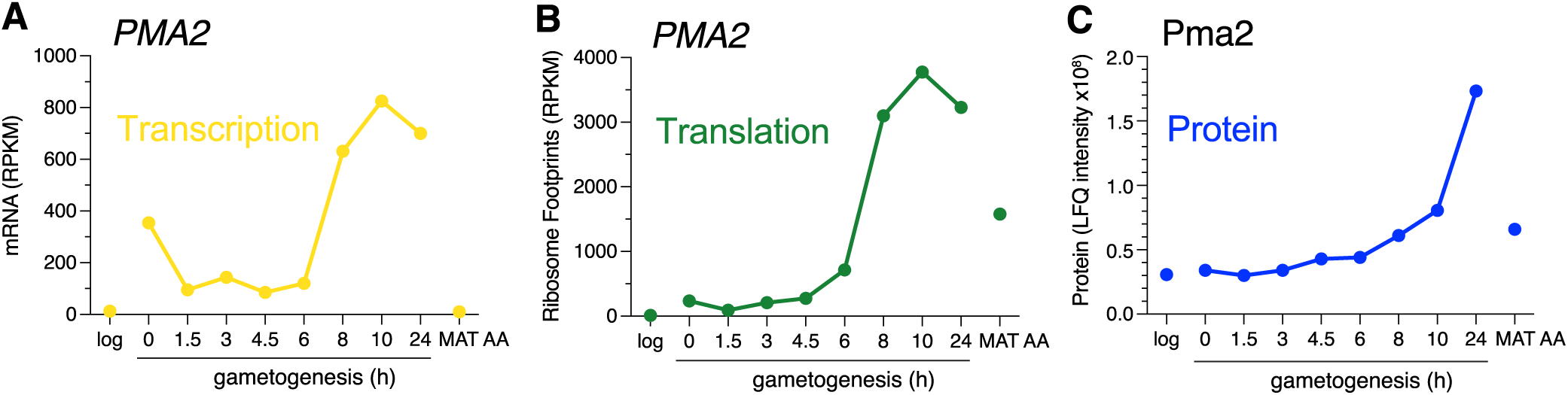
*PMA2* translation and protein levels are upregulated during late gametogenesis. Expression of *PMA2* either during log-phase vegetative growth in 2% glucose containing media (log), the indicated time in gametogenesis inducing media (h), or in control diploids that cannot undergo gametogenesis (*MAT***a**/**a**) plotted from previously generated meiotic datasets (Cheng et al., 2018). (**A**) *PMA2* transcript levels (RNA-seq). (**B**) *PMA2* translation levels (ribosome-protected footprints). (**C**) Pma2 protein levels (mass spectrometry).

**Figure S2.**
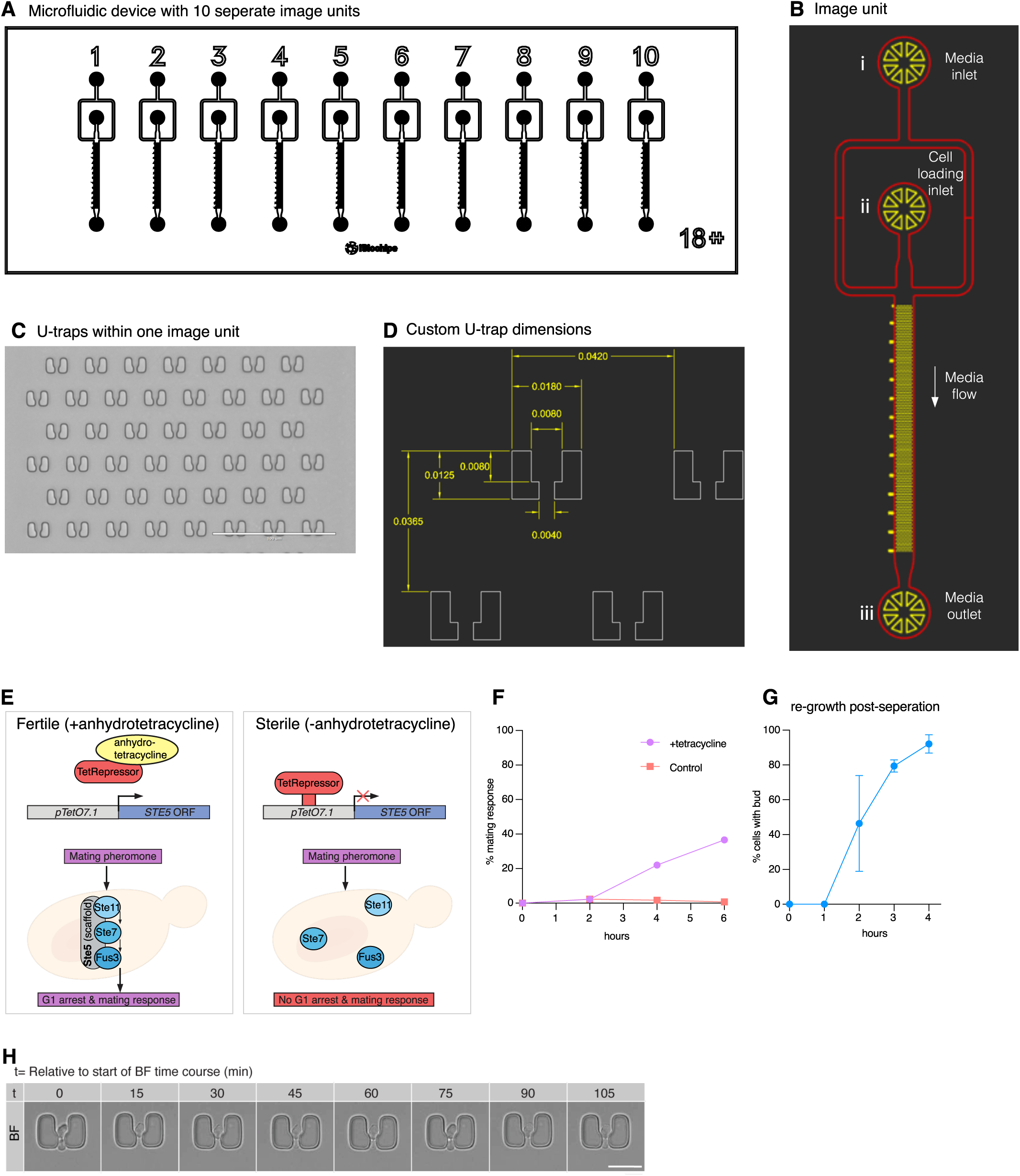
Custom-built U-trap microfluidic devices with specific dimensions for vegetatively growing haploid SK1 cells. (**A**) Schematic view of one microfluidic device with 10 separate imaging units. (**B**) Schematic overview of one unit with (i) media inlet; cell loading inlet (ii) and waste outlet (iii). During the experiment media flows from media inlet (i) to media outlet (iii) while cell loading inlet is plugged with a solid stainless-steel pin. (**C**) Image of U-traps and spacing in one unit, scale bar is 100 µm. (**D**) Custom developed SK1 specific dimensions of trap width and height (8 µm), exit channel width (4 µm), and spacing between traps (24 µm). (**E**) Schematic depiction of genetic system to achieve constitutive sterility. (**F**) Quantification of mating response scoring zygote formation upon 1:1 mixing of *MAT***a** and *MAT***α** cells. (**G**) Monitoring of re-growth of separated gametes in glucose containing liquid culture by scoring emergence of buds. (**H**) Representative trapped gamete dividing twice over the time of two hours, scale bar is 5 µm.

**Figure S3.**
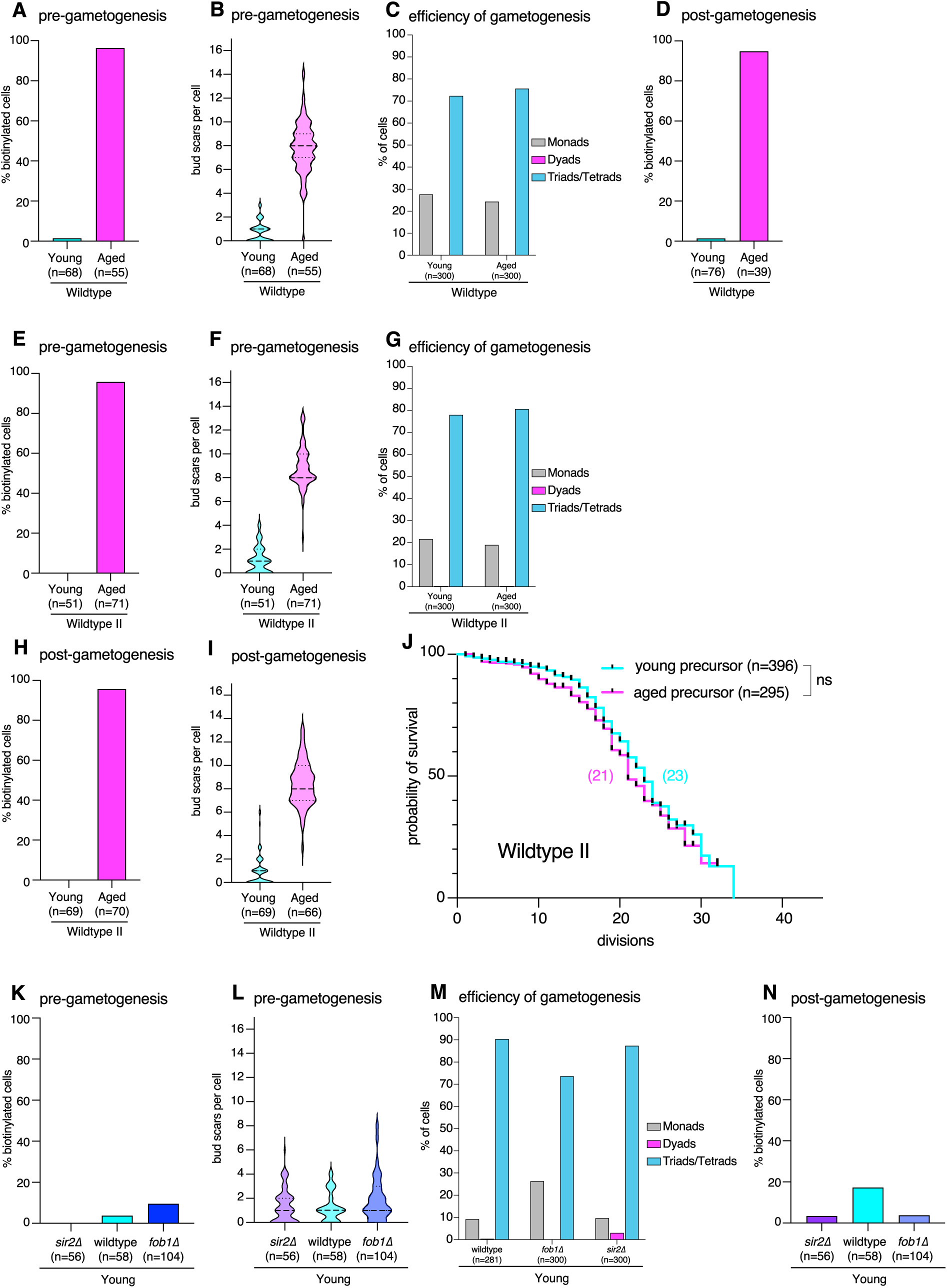
Wildtype gametes completely reset replicative lifespan during gametogenesis. (**A**) Fraction of biotinylated cells in young (1.5%, 95% CI: 0.3–7.9%, n=68) and aged (96.4%, 95% CI: 87.7–99.0%, n=55) wildtype cell populations. (**B**) Quantification of replicative age by assessing number of bud scars in young (0.72± 0.84, mean ± SD, n=68) and aged (7.8 ± 2.2, mean ± SD, n=55) wildtype cell populations. (**C**) Fraction of tetrads in young (72.3%, 95% CI: 67.0–77.1%, n=300) and aged (75.7%, 95% CI: 70.5–80.2% n=300) post-gametogenic cell populations. (**D**) Fraction of biotinylated tetrads in young (1.3%, 95% CI: 0.2–7.1%, n=76) and aged (94.9%, 95% CI: 83.1–98.6%, n=39) wildtype populations after completion of gametogenesis. (**E**) Fraction of biotinylated cells in young (none, n=51) and aged (95.8%, 95% CI: 88.3–98.6%, n=71) wildtype cell populations. (**F**) Quantification of replicative age by assessing number of bud scars in young (1.04± 1.1, mean ± SD, n=51) and aged (8.8 ± 1.7, mean ± SD, n=71) wildtype cell populations. (**G**) Fraction of tetrads in young (78%, 95% CI: 73.0–82.3%, n=300) and aged (80.7%, 95% CI: 75.8–84.7% n=300) post-gametogenic cell populations. (**H**) Fraction of biotinylated tetrads in young (none, n=76) and aged (95.7%, 95% CI: 88.1–98.5%, n=70) wildtype II populations after completion of gametogenesis. (**I**) Quantification of replicative age by assessing number of bud scars in young (0.9± 1.3, mean ± SD, n=69) and aged (8.4± 1.9, mean ± SD, n=66) wildtype II gamete populations. (**J**) Quantification of replicative lifespan of young (median lifespan=23, n=396, 274 censored subjects) and aged (median lifespan=21, n=295, 205 censored subjects, p=0.1341 compared to young) wildtype II gametes. (**K**) Fraction of biotinylated cells in young *sir2Δ* (none, n=78), wildtype (3.7%, 95% CI: 1–12.5%, n=54) and *fob1Δ* (9.5%, 95% CI: 3.8– 22.1%, n=42) cell populations. (**L**) Quantification of replicative age by assessing number of bud scars in young *sir2Δ* (1.3± 1.4, mean ± SD, n=79), wildtype (1.2 ± 1.2, mean ± SD, n=54) and *fob1Δ* (1.9 ± 1.8, mean ± SD, n=42) cell populations. (**M**) Fraction of tetrads in young *sir2Δ* (87.3 %, 95% CI: 83.1– 90.6%, n=300), wildtype (90.4 %, 95% CI: 86.4% – 93.3%, n=281) and *fob1Δ* (73.7%, 95% CI: 68.4–78.3%, n=300) post-gametogenic cell populations. (**N**) Fraction of biotinylated tetrads in young *sir2Δ* (3.4%, 95% CI: 1.0–11.7%, n=58), wildtype (17.3%, 95% CI: 11.2–25.7%, n=104) and *fob1Δ* (3.8%, 95% CI: 1.0–12.8%, n=53) populations after completion of gametogenesis. Confidence intervals were calculated using the Wilson score method and statistical significance for differences between replicative lifespans was calculated using the Log-rank (Mantel-cox) test, tick marks on survival curves represent censored events.

**Figure S4.**
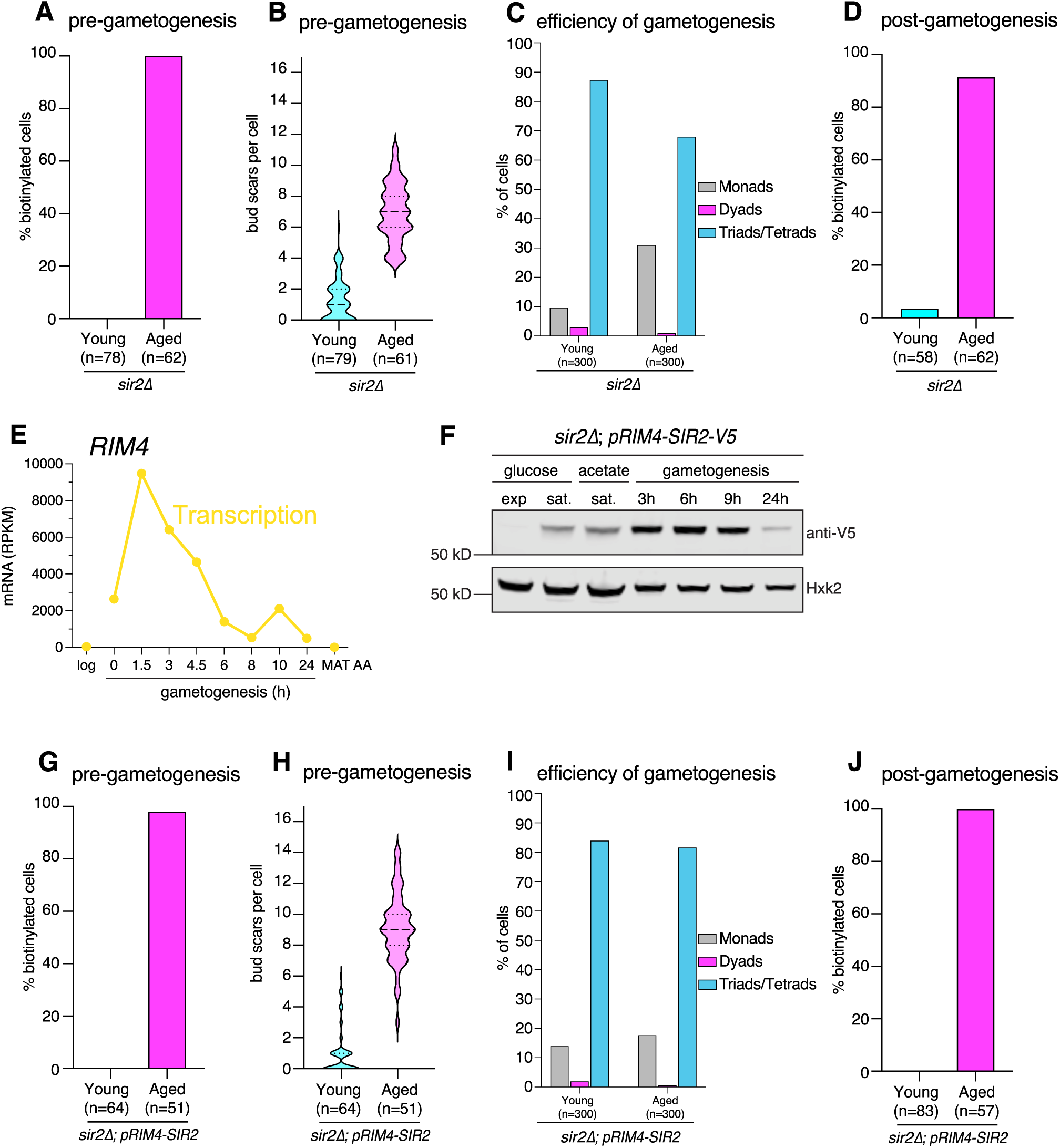
*sir2Δ* mutants and gametogenesis specific Sir2 expression both fail to reset replicative lifespan of gametes. (**A**) Fraction of biotinylated cells in young (none, n=78, same data as shown in Fig. 4A) and aged (all, n=78) *sir2Δ* cell populations. (**B**) Quantification of replicative age by assessing number of bud scars in young (1.3± 1.4, mean ± SD, n=79, same data as shown in Fig. 4B) and aged (7± 1.9, mean ± SD, n=61) *sir2Δ* cell populations. (**C**) Fraction of tetrads in young (87.3% 95% CI: 83.1–90.6% n=300, same data as shown in Fig. 4E) and aged (68.0% (95% CI: 62.5–73.0%, n=300) post-gametogenic *sir2Δ* cell populations. (**D**) Fraction of biotinylated tetrads in young (3.4%, 95% CI: 1.0–11.7%, n=58, same data as shown in Fig. 4C) and aged (91.9%, 95% CI: 82.5–96.5%, n=62) *sir2Δ* populations after completion of gametogenesis. (**E**) *RIM4* transcript levels (RNA-seq) either during log-phase vegetative growth in 2% glucose containing media (log), the indicated time in gametogenesis inducing media (h), or in control diploids that cannot undergo gametogenesis (*MAT***a**/**a**) plotted from previously generated meiotic datasets (Cheng et al., 2018). (**F**) Immunoblot to quantify Sir2-V5 levels driven by *RIM4* promoter fragment (anti-V5) and loading control (Hxk2) in growth media containing ether 2% glucose (glucose) or acetate (acetate) and at different timepoints upon transfer to gametogenesis-inducing media (gametogenesis). (**G**) Fraction of biotinylated cells in young (none, n=64) and aged (98.0%, 95% CI: 89.7–99.7%, n=51) *pRIM4-SIR2 sir2Δ* cell populations. (**H**) Quantification of replicative age by assessing number of bud scars in young (0.98± 1.5, mean ± SD, n=64) and aged (9.1± 2.2 (mean ± SD, n=51) *pRIM4-SIR2 sir2Δ* cell populations. (**I**) Fraction of tetrads in young (84 %, 95% CI: 79.4–87.7%, n=300) and aged (81.7%, 95% CI: 76.9–85.6%, n=300) post-gametogenic *pRIM4-SIR2 sir2Δ* cell populations. (**J**) Fraction of biotinylated tetrads in young (none, n=83) and aged (all, n=57) *pRIM4-SIR2 sir2Δ* populations after completion of gametogenesis. Confidence intervals were calculated using the Wilson score method.

**Figure S5.**
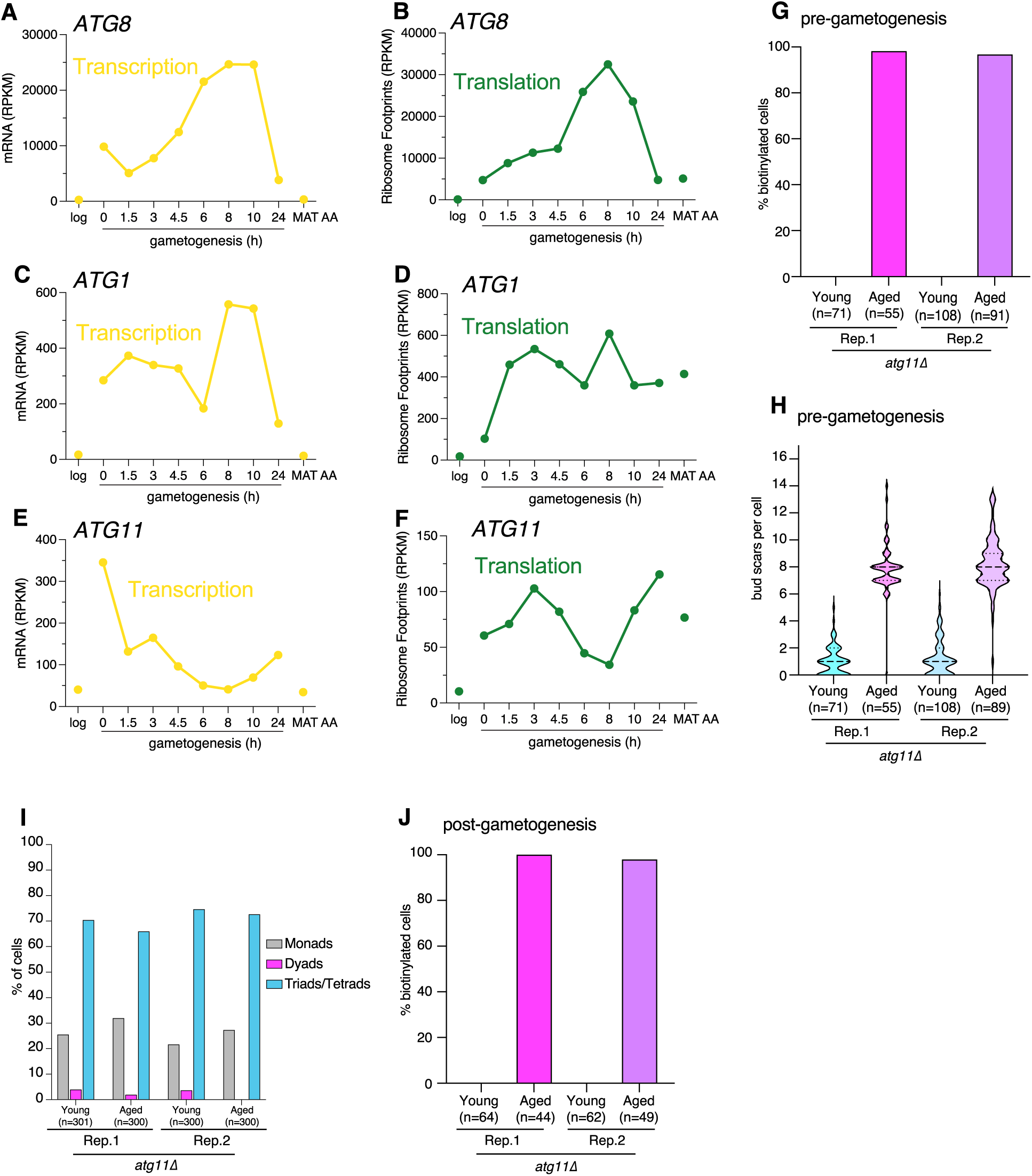
*atg11Δ* mutants fail to reset replicative lifespan of gametes. (**A-F**) Expression of autophagy factors either during log-phase vegetative growth in 2% glucose containing media (log), the indicated time in gametogenesis inducing media (h), or in control diploids that cannot undergo gametogenesis (*MAT***a**/**a**) plotted from previously generated meiotic datasets (Cheng et al., 2018). (**A**) *ATG8* transcript levels (RNA-seq). (**B**) *ATG8* translation levels (ribosome-protected footprints). (**C**) *ATG1* transcript levels (RNA-seq). (**D**) *ATG1* translation levels (ribosome-protected footprints). (**E**) *ATG11* transcript levels (RNA-seq). (**F**) *ATG11* translation levels (ribosome-protected footprints). (**G**) Fraction of biotinylated cells in *atg11Δ* cell populations: Replicate 1 (Rep. 1) young (none, n=71) and aged (98.2%, 95% CI: 90.4–99.7%, n=55); Replicate 2 (Rep. 2) young (none, n=108) and aged (96.7%, 95% CI: 90.8–98.9%, n=91). (**H**) Quantification of replicative age by assessing number of bud scars in *atg11Δ* cell populations: Rep. 1 young (1 ± 1.1, mean ± SD, n=71) and aged (7.9± 1.9, mean ± SD, n=55); Rep. 2 young (1.2 ± 1.3, mean ± SD, n=108) and aged (8.2± 1.9, mean ± SD, n=89). (**I**) Fraction of tetrads in post-gametogenic *atg11Δ* cell populations: Rep.1 young (70.4%, 95% CI: 65.0–75.3%), n=301) and aged (66.0%, 95% CI: 60.5–71.1%, n=300); Rep. 2 young (74.7%, 95% CI: 69.5–79.3%, n=300) and aged (72.7%, 95% CI: 67.4–77.4%, n=300). (**J**) Fraction of biotinylated tetrads in *atg11Δ* populations after completion of gametogenesis: Rep.1 young (none, n=64) and aged (all, n=44; Rep. 2 young (none, n=62) and aged (98%, 95% CI: 89.3–99.6%, n=91all, n=49). Confidence intervals were calculated using the Wilson score method.

**Figure S6.**
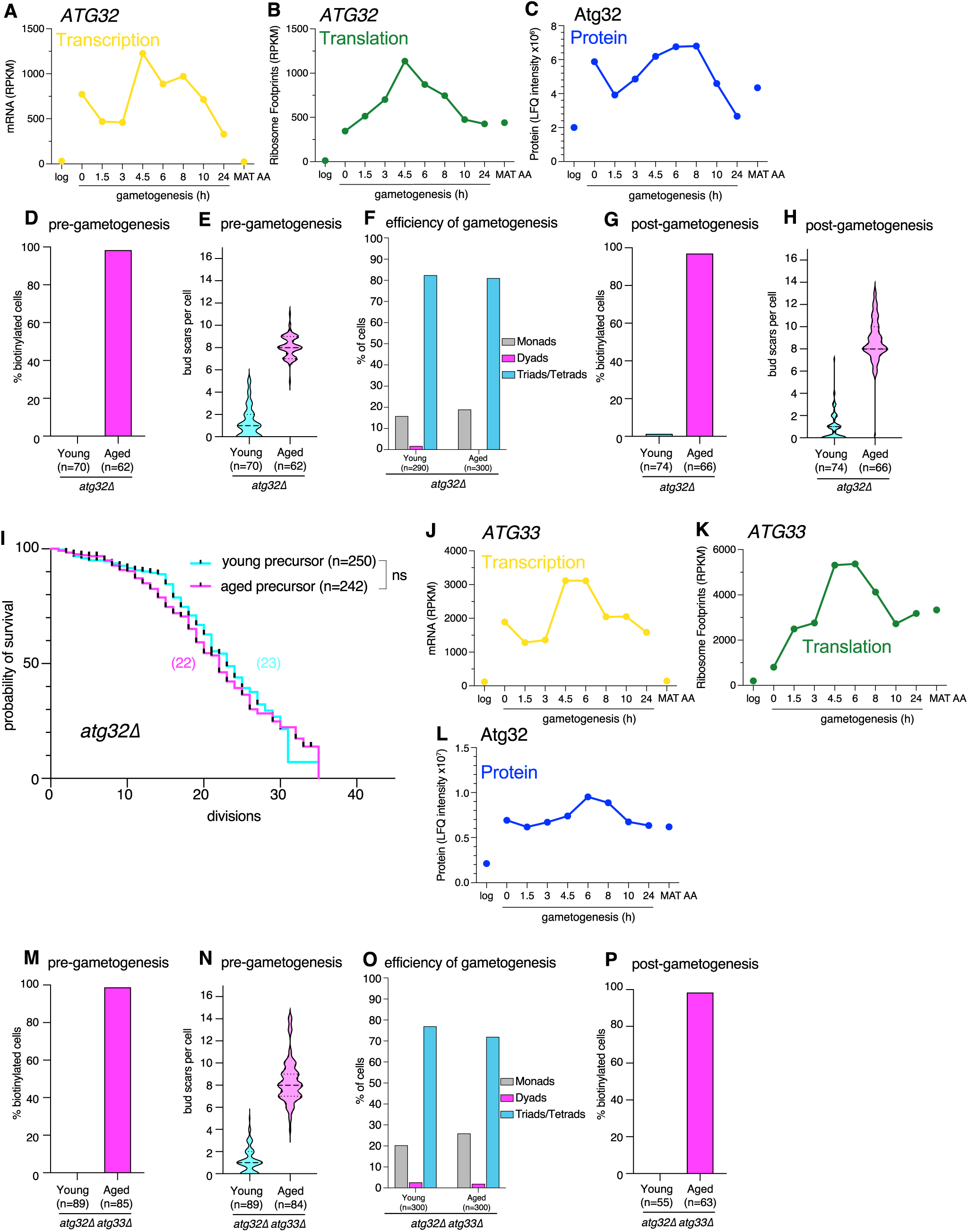
*atg32Δ* mutants and *atg32Δ atg33Δ* double mutants completely reset replicative lifespan of gametes. (**A-C & J-L**) Expression levels either during log-phase vegetative growth in 2% glucose containing media (log), the indicated time in gametogenesis inducing media (h), or in control diploids that cannot undergo gametogenesis (*MAT***a**/**a**) plotted from previously generated meiotic datasets (Cheng et al., 2018). (**A**) *ATG32* transcript levels (RNA-seq). (**B**) *ATG32* translation levels (ribosome-protected footprints). (**C**) Atg32 protein levels (mass spectrometry). (**D**) Fraction of biotinylated cells in young (none, n=70) and aged (98.4%, 95% CI: 91.4–99.7%, n=62) *atg32Δ* cell populations. (**E**) Quantification of replicative age by assessing number of bud scars in young (1.3± 1.4, mean ± SD, n=70) and aged (8.1± 0.98, mean ± SD, n=62) *atg32Δ* cell populations. (**F**) Fraction of tetrads in young (82.4%, 95% CI: 77.6–86.4%), n=290) and aged (81.0%, 95% CI: 76.2–85.0%, n=300) post-gametogenic *atg32Δ* cell populations. (**G**) Fraction of biotinylated tetrads in young (1.4%, 95% CI: 0.2–7.3%, n=74) and aged 97.0% (95% CI: 89.6–99.2%, n=66) *atg32Δ* populations after completion of gametogenesis. (**H**) Quantification of replicative age by assessing number of bud scars in young (0.93± 1.3, mean ± SD, n=74) and aged (8.7± 2, mean ± SD, n=66) *atg32Δ* gamete populations. (**I**) Quantification of replicative lifespan of young (median lifespan=23, n=242, 151 censored subjects, 1 technical replicate) and aged (median lifespan=22, n=250, 151 censored subjects, p=0.2691) *atg32Δ* gametes. (**J**) *ATG33* transcript levels (RNA-seq). (**K**) *ATG33* translation levels (ribosome-protected footprints). (**L**) Atg33 protein levels (mass spectrometry). (**M**) Fraction of biotinylated cells in young (none, n=89) and aged (98.8%, 95% CI: 93.6–99.8%, n=85) *atg32Δ atg33Δ* double mutant cell populations. (**N**) Quantification of replicative age by assessing number of bud scars in young (1.2± 1.1, mean ± SD, n=89) and aged (8.4± 2, mean ± SD, n=84) *atg32Δ atg33Δ* double mutants. (**O**) Fraction of tetrads in young (77.0%, 95% CI: 71.9–81.4%, n=300) and aged (72.0%, 95% CI: 66.7–76.8%, n=300) post-gametogenic *atg32Δ atg33Δ* cell populations. (**P**) Fraction of biotinylated tetrads in young (none, n=55) and aged (98.4%, 95% CI: 91.5–99.7%, n=63) *atg32Δ atg33Δ* populations after completion of gametogenesis. Confidence intervals were calculated using the Wilson score method and statistical significance for differences between replicative lifespans was calculated using the Log-rank (Mantel-cox) test, tick marks on survival curves represent censored events.

**Figure S7.**
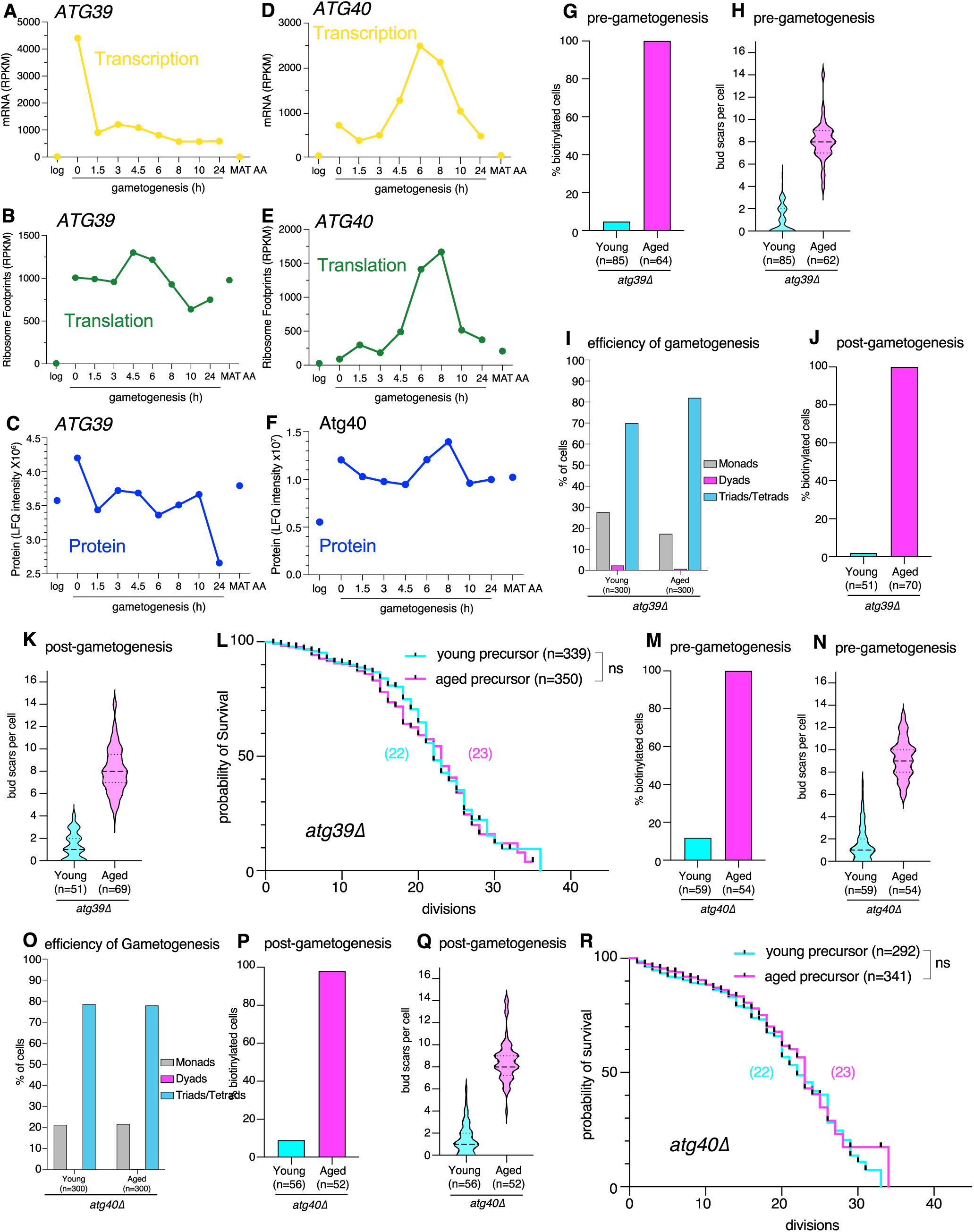
*atg39Δ* and *atg40Δ* single mutants reset replicative lifespan during gametogenesis. (**A-F**) Expression of *ATG39* and *ATG40* either during log-phase vegetative growth in 2% glucose containing media (log), the indicated time in gametogenesis inducing media (h), or in control diploids that cannot undergo gametogenesis (*MAT***a**/**a**) plotted from previously generated meiotic datasets (Cheng et al., 2018). (**A**) *ATG39* transcript levels (RNA-seq). (**B**) *ATG39* translation levels (ribosome-protected footprints). (**C**) Atg39 protein levels (mass spectrometry). (**D**) *ATG40* transcript levels (RNA-seq). (**E**) *ATG40* translation levels (ribosome-protected footprints). (**F**) Atg40 protein levels (mass spectrometry). (**G**) Fraction of biotinylated cells in young (3.5%, 95% CI: 1.2–9.9%, n=85) and aged (all, n=64) *atg39Δ* cell populations. (**H**) Quantification of replicative age by assessing number of bud scars in young (0.98± 1.2, mean ± SD, n=85) and aged (8.1± 1.8, mean ± SD, n=64) *atg39Δ* cell populations. (**I**) Fraction of tetrads in young (70.0%, 95% CI: 64.6–74.9%, n=300) and aged (82.0% (95% CI: 77.3–85.9%, n=300) post-gametogenic *atg39Δ* cell populations. (**J**) Fraction of biotinylated tetrads in young (2.0%, 95% CI: 0.3–10.3%, n=51) and aged (all, n=69) *atg39Δ* populations after completion of gametogenesis. (**K**) Quantification of replicative age by assessing number of bud scars in young (1.3± 1.2, mean ± SD, n=51) and aged (8.1± 2.1, mean ± SD, n=69) *atg39Δ* gamete populations. (**L**) Quantification of replicative lifespan of young (median lifespan=22, n=339, 188 censored subjects) and aged *atg39Δ* (median lifespan=23, n=350, 225 censored subjects, p=0.418 compared to young) gametes. (**M**) Fraction of biotinylated cells in young (11.9%, 95% CI: 5.9–22.5%, n=59) and aged (all, n=54) *atg40Δ* cell populations. (**N**) Quantification of replicative age by assessing number of bud scars in young (1.6± 1.6, mean ± SD, n=59) and aged (9.3± 1.8, mean ± SD, n=54) *atg40Δ* cell populations. (**O**) Fraction of tetrads in young (78.7%, 95% CI: 73.7–82.9%, n=300) and aged (78.0%, 95% CI: 73.0–82.3%, n=300) post-gametogenic *atg39Δ* cell populations. (**P**) Fraction of biotinylated tetrads in young (8.9%, 95% CI: 3.9–19.3%, n=56) and aged (98.1%, 95% CI: 89.9–99.7%, n=52) *atg40Δ* populations after completion of gametogenesis. (**Q**) Quantification of replicative age by assessing number of bud scars in young (1.4± 1.4, mean ± SD, n=56), and aged (8.6± 1.9, mean ± SD, n=52) *atg40Δ* gamete populations. (**R**) Quantification of replicative lifespan of young (median lifespan=22, n=292, 176 censored subjects) and aged *atg40Δ* (median lifespan=23, n=341, 256 censored subjects, p=0.389 compared to young) gametes. Confidence intervals were calculated using the Wilson score method and statistical significance for differences between replicative lifespans was calculated using the Log-rank (Mantel-cox) test, tick marks on survival curves represent censored events.

**Figure S8.**
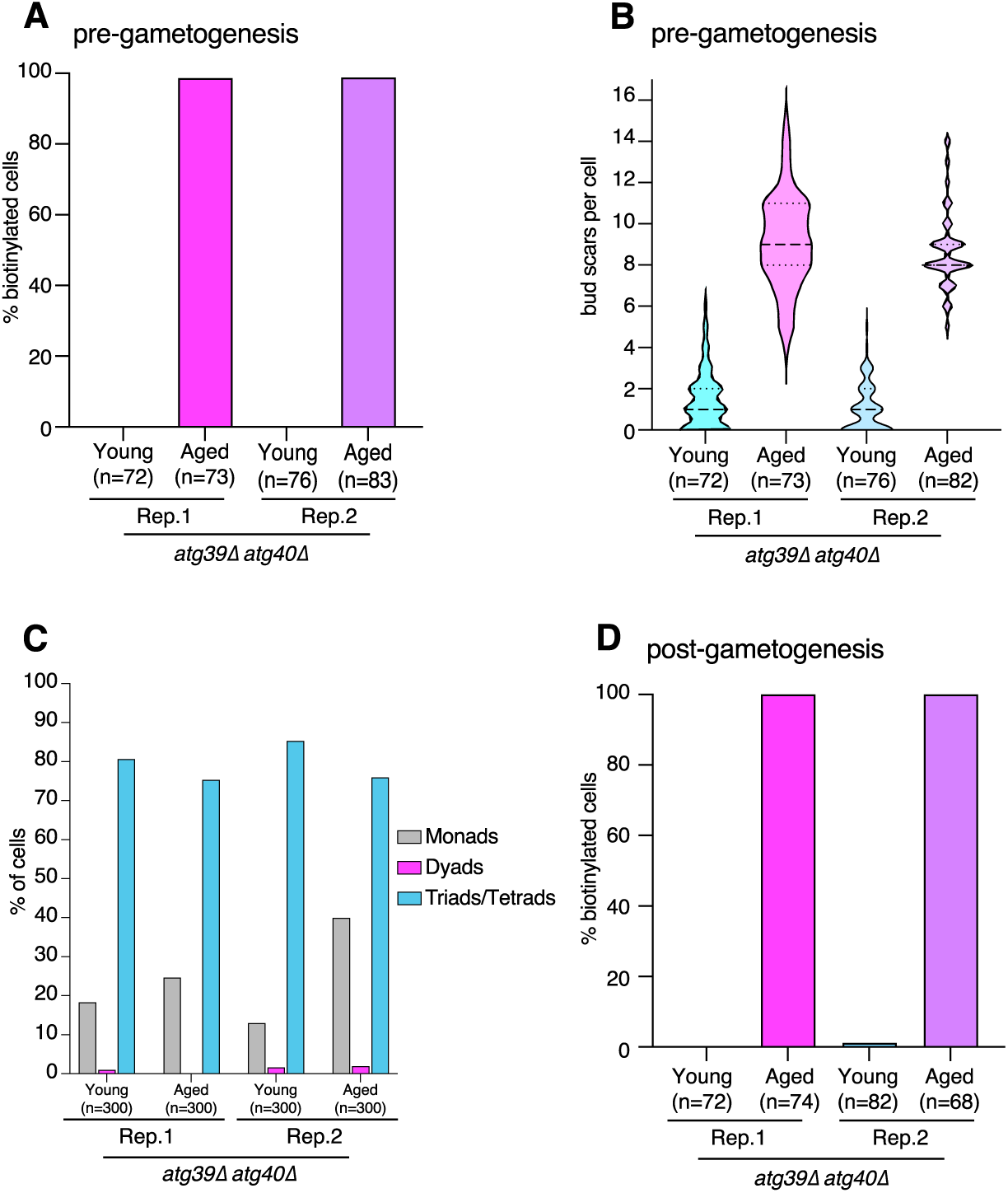
*atg39Δ atg40Δ* double mutants fail to reset replicative lifespan during gametogenesis. (**A**) Fraction of biotinylated cells in *atg39Δ atg40Δ* cell populations: Replicate 1 (Rep. 1) young (none, n=72) and aged (98.6%, 95% CI: 92.6–99.8%, n=73); Replicate 2 (Rep. 2) young (none, n=76) and aged (98.8%, 95% CI: 93.5–99.8%, n=83). (**B**) Quantification of replicative age by assessing number of bud scars in *atg39Δ atg40Δ* cell populations: Rep. 1 young (1.4± 1.4, mean ± SD, n=72) and aged (9.1± 2.3, mean ± SD, n=73); Rep. 2 young (1 ± 1.2, mean ± SD, n=76) and aged (8.7± 1.9, mean ± SD, n=82). (**C**) Fraction of tetrads in post-gametogenic *atg39Δ atg40Δ* cell populations: Rep.1 young (80.7%, 95% CI: 75.8–84.7%, n=300) and aged (75.3%, 95% CI: 70.2–79.9%, n=300); Rep. 2 young (85.3%, 95% CI: 80.9–88.9%, n=300) and aged (76.0%, 95% CI: 70.9–80.5%, n=300). (**D**) Fraction of biotinylated tetrads in *atg39Δ atg40Δ* populations after completion of gametogenesis: Rep.1 young (none, n=72) and aged (all, n=74); Rep. 2 young (1.2%, 95% CI: 0.2–6.6%, n=82) and aged (all, n=68). Confidence intervals were calculated using the Wilson score method.

**Figure S9.**
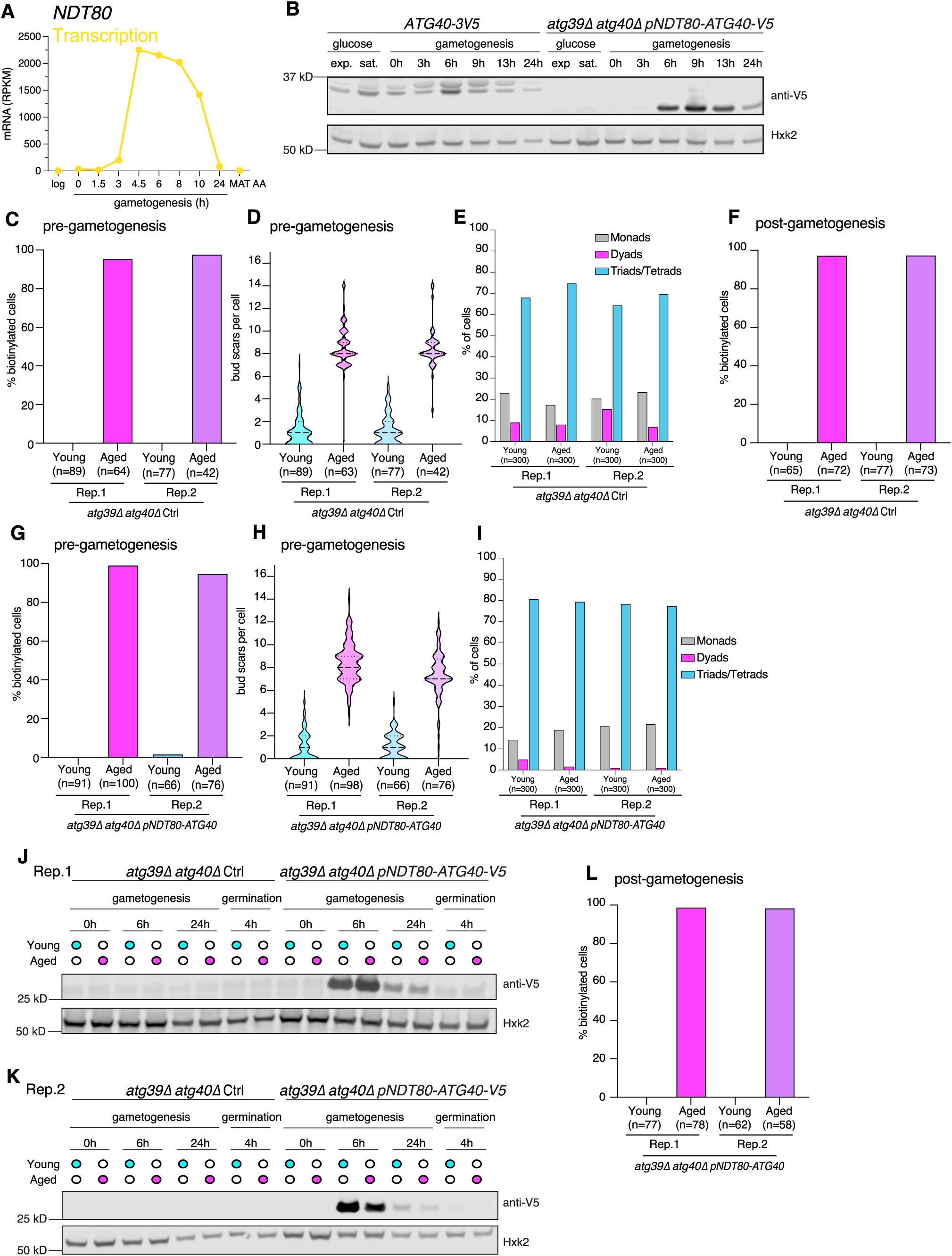
Supporting data for meiosis specific expression of *ATG40* is sufficient to rescue reset of replicative lifespan in cells that lack *ATG40 and ATG39*. **A**) *NDT80* transcript levels (RNA-seq) either during log-phase vegetative growth in 2% glucose containing media (log), the indicated time in gametogenesis inducing media (h), or in control diploids that cannot undergo gametogenesis (*MAT***a**/**a**) plotted from previously generated meiotic datasets (Cheng et al., 2018). (**B**) Immunoblot comparing endogenous Atg40-3V5 to pNDT80-Atg40-V5 rescue construct protein levels (anti-V5) and loading control (Hxk2) during exponential and saturated growth in media containing 2% glucose (glucose) and at different timepoints upon transfer to gametogenesis-inducing media (gametogenesis). (**C**) Fraction of biotinylated cells in *atg39Δ atg40Δ* control cell populations: Replicate 1 (Rep. 1) young (none, n=89) and aged (95.3%, 95% CI: 87.1–98.4%, n=64); Replicate 2 (Rep. 2) young (none, n=77) and aged (97.6%, 95% CI: 87.7–99.6%, n=44). (**D**) Quantification of replicative age by assessing number of bud scars in *atg39Δ atg40Δ* control cell populations: Rep. 1 young (1.2± 1.4, mean ± SD, n=89) and aged (8.5± 1.9, mean ± SD, n=63); Rep. 2 young (1.3 ± 1.3, mean ± SD, n=77) and aged (8.5± 1.9, mean ± SD, n=42). (**E**) Fraction of tetrads in post-gametogenic *atg39Δ atg40Δ* control cell populations: Rep.1 young (68.0%, 95% CI: 62.5–73.0%, n=300) and aged (74.7%, 95% CI: 69.5–79.3%, n=300); Rep. 2 young (64.3%, 95% CI: 58.8–69.5%, n=300) and aged (69.7%, 95% CI: 64.2–74.6%, n=300). (**F**) Fraction of biotinylated tetrads in *atg39Δ atg40Δ* control populations after completion of gametogenesis: Rep.1 young (none, n=65) and aged (97.2%, 95% CI: 90.4–99.2%, n=72); Rep. 2 young (none, n=77) and aged (97.3%, 95% CI: 90.5–99.2%, n=73). (**G**) Fraction of biotinylated cells in *atg39Δ atg40Δ pNDT80-ATG40* cell populations: Replicate 1 (Rep. 1) young (none, n=91) and aged (99.0%, 95% CI: 94.6–99.8%, n=100); Replicate 2 (Rep. 2) young (1.5%, 95% CI: 0.3–8.1%, n=66) and aged (94.7%, 95% CI: 87.2–97.9%, n=76). (**H**) Quantification of replicative age by assessing number of bud scars in *atg39Δ atg40Δ pNDT80-ATG40* cell populations: Rep. 1 young (1.0± 1.3, mean ± SD, n=91) and aged (8.4± 1.3, mean ± SD, n=98); Rep. 2 young (1.1 ± 1.1, mean ± SD, n=66) and aged (7.3± 2.1, mean ± SD, n=76). (**I**) Fraction of tetrads in post-gametogenic *atg39Δ atg40Δ pNDT80-ATG40* cell populations: Rep.1 young (80.7%, 95% CI: 75.8–84.7%, n=300) and aged (79.3%, 95% CI: 74.4–83.5%, n=300); Rep. 2 young (78.3%, 95% CI: 73.3–82.6%, n=300) and aged (77.3%, 95% CI: 72.3–81.7%, n=300). (**J&K**) Immunoblot of young and aged control and pNDT80-Atg40-V5 samples to quantify protein levels (anti-V5) and loading control (Hxk2) at different timepoints upon transfer to gametogenesis-inducing media (gametogenesis) and after re-growth in media containing 2% glucose (germination). (**J**) Samples collected from Rep.1. **(K**) Samples collected from Rep.2. (**L**) Fraction of biotinylated tetrads in *atg39Δ atg40Δ pNDT80-ATG40* cell populations after completion of gametogenesis: Rep.1 young (none, n=77) and aged (98.7%, 95% CI: 93.1–99.8%, n=78); Rep. 2 young (none, n=62) and aged (98.3%, 95% CI: 90.9–99.7%, n=58). Confidence intervals were calculated using the Wilson score method.

**Figure S10.**
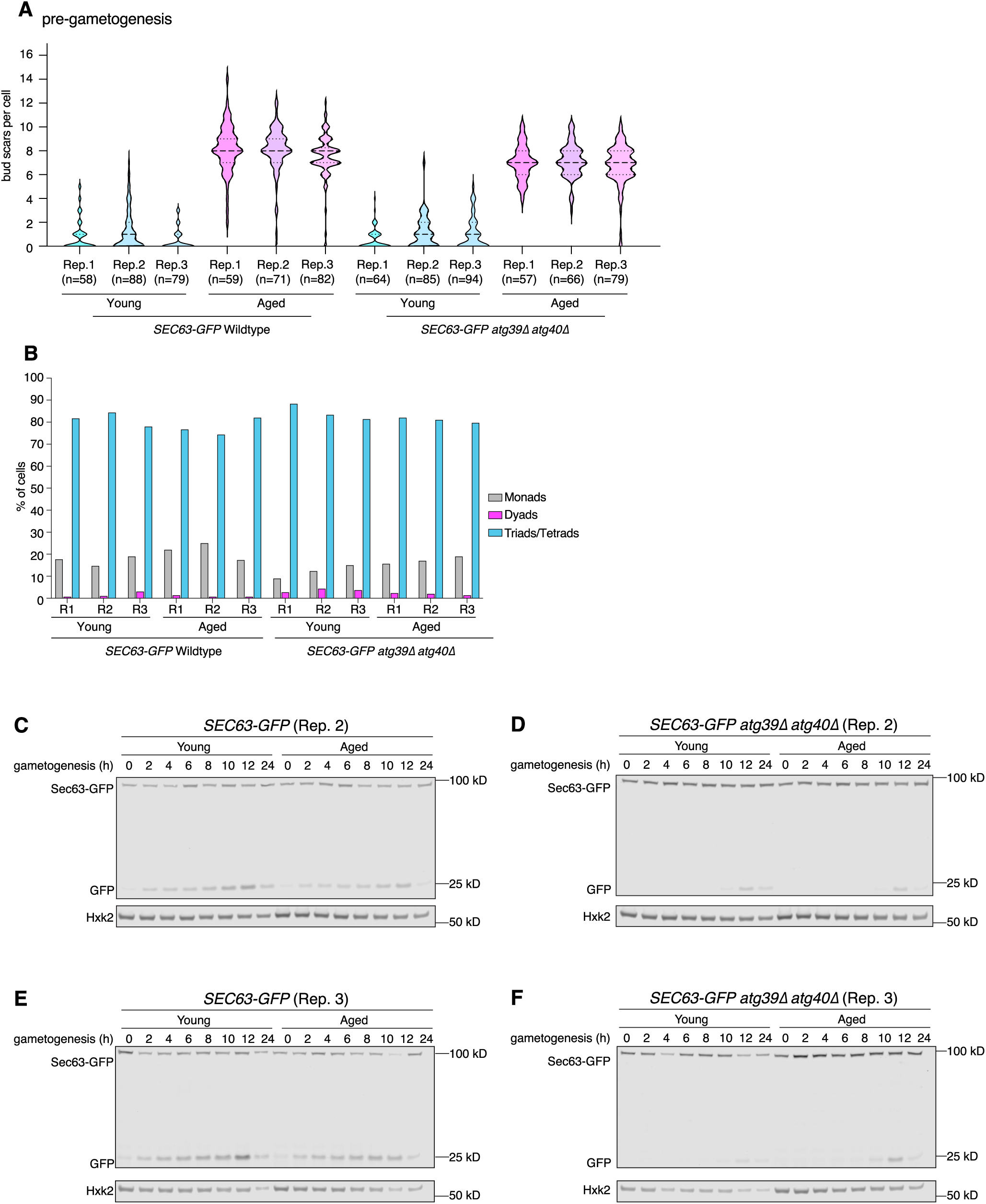
Supporting data for accumulation of ER components during gametogenesis in cells that lack *ATG40 and ATG39*. (**A**) Quantification of replicative age by assessing number of bud scars in *SEC63-GFP* and *SEC63-GFP atg39Δ atg40Δ* cell populations. *SEC63-GFP* cell populations: Rep. 1 young (0.9± 1.3, mean ± SD, n=58) and aged (8.2± 2, mean ± SD, n=59); Rep. 2 young (1.3 ± 1.7, mean ± SD, n=88) and aged (8± 2, mean ± SD, n=71)); Rep. 3 young (0.51 ± .89, mean ± SD, n=79) and aged (7.5± 2, mean ± SD, n=82). *SEC63-GFP atg39Δ atg40Δ* cell populations: Rep. 1 young (0.55± 0.85, mean ± SD, n=64) and aged (7± 1.4, mean ± SD, n=57); Rep. 2 young (1.1 ± 1.4, mean ± SD, n=85) and aged (7.2± 1.4, mean ± SD, n=66); Rep. 2 young (1.1 ± 1.4, mean ± SD, n=94) and aged (6.7± 1.9, mean ± SD, n=79). (**B**) Fraction of tetrads in post gametogenic *SEC63-GFP* and *SEC63-GFP atg39Δ atg40Δ* cell populations. *SEC63-GFP* cell populations: Rep.1 young (81.7%, 95% CI: 76.9–85.6%, n=300) and aged (76.7%, 95% CI: 71.6–81.1%, n=300); Rep. 2 young (84.3%, 95% CI: 79.83–88.0%, n=300) and aged (74.3%, 95% CI: 69.1–78.9%, n=300); Rep. 3 young (78.0%, 95% CI: 73.0–82.3%, n=300) and aged (82.0%, 95% CI: 77.3–85.9%, n=300). *SEC63-GFP atg39Δ atg40Δ* cell populations: Rep.1 young (88.3%, 95% CI: 84.2–91.5%, n=300) and aged (82.0%, 95% CI: 77.3–85.9%, n=300); Rep. 2 young (83.3%, 95% CI: 78.7–87.1%, n=300) and aged (81.0%, 95% CI: 76.2–85.0%, n=300); Rep. 3 young (81.3%, 95% CI: 76.5–85.3%, n=300) and aged (79.7%, 95% CI: 74.8–83.8%, n=300). **C)** Immunoblot of protein samples form young and aged *SEC63-GFP* cells to quantify protein levels (anti-GFP) and loading control (Hxk2) at different timepoints upon transfer to gametogenesis-inducing media (gametogenesis (h)), second replica used for quantification of Sec63-GFP levels normalize to loading control (Hxk2) (Fig. 6E). **D)** Immunoblot of protein samples form young and aged *atg39Δ atg40Δ SEC63-GFP* cells to quantify protein levels (anti-GFP) and loading control (Hxk2) at different timepoints upon transfer to gametogenesis-inducing media (gametogenesis (h)), second replica used for quantification of Sec63-GFP levels normalize to loading control (Hxk2) (Fig. 6E). **E)** Immunoblot of protein samples form young and aged *SEC63-GFP* cells to quantify protein levels (anti-GFP) and loading control (Hxk2) at different timepoints upon transfer to gametogenesis-inducing media (gametogenesis (h)), third replica used for quantification of Sec63-GFP levels normalize to loading control (Hxk2) (Fig. 6E). **F)** Immunoblot of protein samples form young and aged *atg39Δ atg40Δ SEC63-GFP* cells to quantify protein levels (anti-GFP) and loading control (Hxk2) at different timepoints upon transfer to gametogenesis-inducing media (gametogenesis (h)), third replica used for quantification of Sec63-GFP levels normalize to loading control (Hxk2) (Fig. 6E).

## Supplementary Tables

**Table S1.**
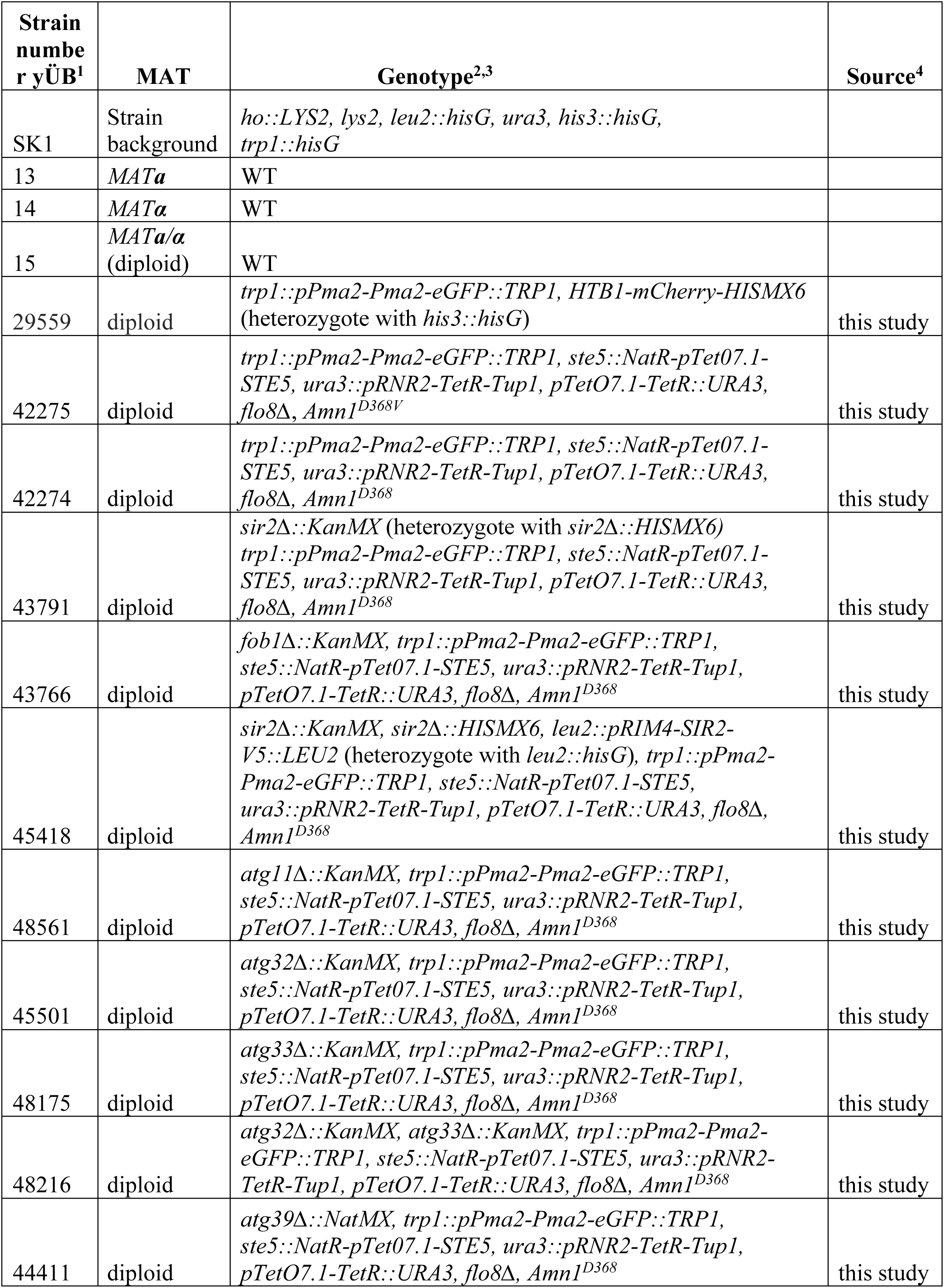
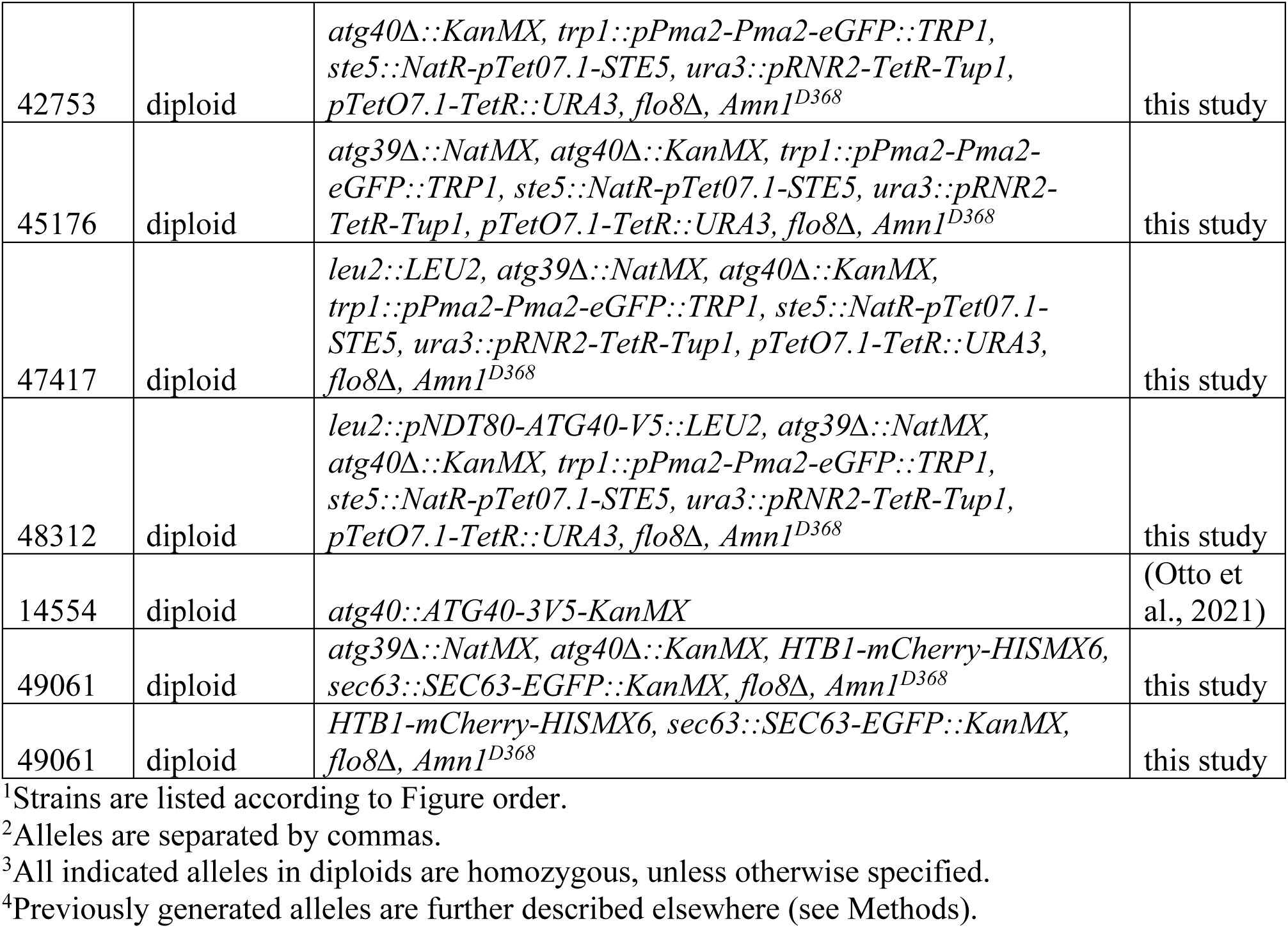
Strain Table.

**Table S2.** Plasmids.

| Plasmid number (pÜB) | Construct<sup>1</sup> | Backbone | Bacterial selection | Source / Reference |
| --- | --- | --- | --- | --- |
| 1 | *KanMX6* | pFA6a | Ampicillin | (Longtine et al., 1998) |
| 153 | *NatMX4* | pFA6a | Ampicillin | (Longtine et al., 1998) |
| 182 | *linker-eGFP_Kan* | pFA6a |  | Adapted from (Longtine et al., 1998) |
| 2693 | *NatMX4_PtetO7.1* | pFA6a | Ampicillin | Adapted from (Azizoglu et al., 2021) |
| 2255 | *kanMX6-ins1* | pFA6a | Ampicillin | (Powers et al., 2022) |
| 2309 | *his3MX6-ins3* | pFA6a | Ampicillin | (Powers et al., 2022) |
| 1905 | *PPMA2-PMA2-eGPF LEU2* | pNH604 | Ampicillin | This study |
| 2991 | *PRIM4-SIR2-V5 LEU2* | pLC605 | Ampicillin | This study |
| 3076 | *PNDT80-ATG40-V5 LEU2* | pÜB2991 | Ampicillin | This study |
| 2371 | *PRNR2-TetR-TUP1 PtetO7.1-TetR URA3* | pSV606 | Ampicillin | Adapted from (Azizoglu et al., 2021) |

<sup>1</sup>Underscores indicate separation between plasmid elements (e.g. construct and selection cassette)

**Table S3.**
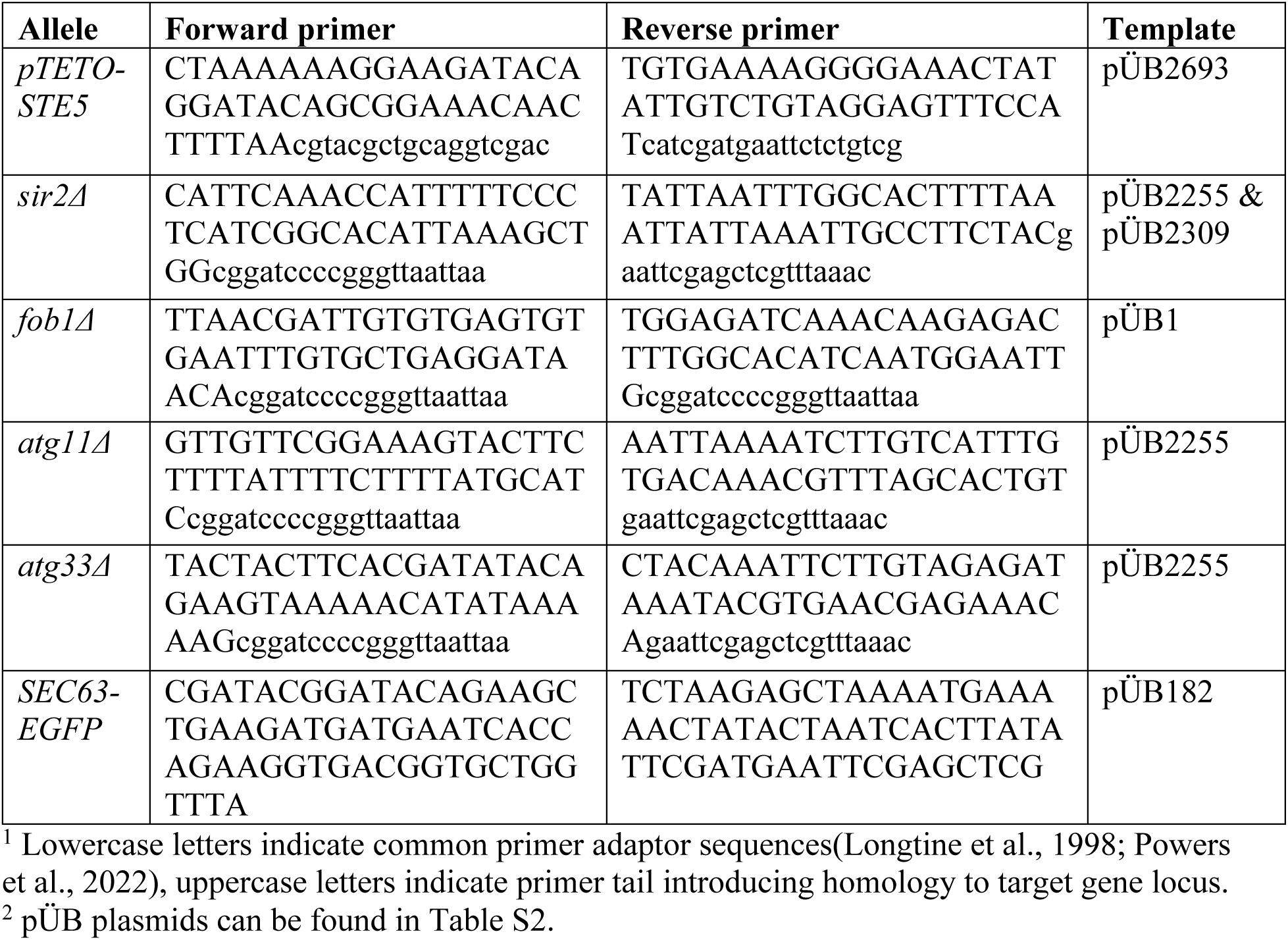
Primers.

